# OGG1 and MUTYH repair activities promote telomeric 8-oxoguanine induced cellular senescence

**DOI:** 10.1101/2023.04.10.536247

**Authors:** Mariarosaria De Rosa, Ryan P. Barnes, Prasanth R. Nyalapatla, Peter Wipf, Patricia L. Opresko

**Affiliations:** Department of Environmental and Occupational Health, University of Pittsburgh School of Public Health, Pittsburgh, PA, USA; UPMC Hillman Cancer Center, Pittsburgh, PA, USA; Deparment of Chemistry, University of Pittsburgh, Pittsburgh, PA, USA; Department of Pharmacology and Chemical Biology, University of Pittsburgh School of Medicine, Pittsburgh, PA, USA

## Abstract

Telomeres are prone to formation of the common oxidative lesion 8-oxoguanine (8oxoG), and the acute production of 8oxoG damage at telomeres is sufficient to drive rapid cellular senescence. OGG1 and MUTYH glycosylases initiate base excision repair (BER) at 8oxoG sites to remove the lesion or prevent mutation. Here, we show OGG1 loss or inhibition, or MUTYH loss, partially rescues telomeric 8oxoG-induced senescence, and loss of both glycosylases results in a near complete rescue. Loss of these glycosylases also suppresses 8oxoG-induced telomere fragility and dysfunction, indicating that single-stranded break (SSB) intermediates arising downstream of glycosylase activity impair telomere replication. The failure to initiate BER in glycosylase-deficient cells suppresses PARylation at SSB intermediates and confers resistance to the synergistic effects of PARP inhibitors on damage-induced senescence. Our studies reveal that inefficient completion of 8oxoG BER at telomeres triggers cellular senescence via SSB intermediates which impair telomere replication and stability.

## INTRODUCTION

Telomeres are nucleoprotein structures that protect the ends of linear chromosomes. In vertebrates, telomeres consist of tandem TTAGGG repeats and end in a single-stranded 3’ G-rich overhang that forms a loop structure in cooperation with shelterin, the telomere capping complex ^1, 2^. Telomeric DNA structure and shelterin prevent the chromosome end from being falsely recognized as a chromosome break ^3^. Telomere shortening during cell division and critically short, dysfunctional telomeres activate the canonical DNA damage response (DDR), leading to p53 activation and downstream effectors that enforce proliferative arrest and senescence, apoptosis or autophagy depending on the cell type or context ^3–5^. A few dysfunctional telomeres are sufficient to trigger senescence in human fibroblasts ^6^. Dysfunctional telomeres, detected by the co-localization of telomeric DNA with DDR factors γH2AX or 53BP1, are hallmarks of senescent cells and increase in various cell types and tissue with age, and in diseases associated with oxidative stress and inflammation ^7–9^. Dysfunctional telomeres have been observed even in the absence of shortening and extensive cell divisions in low proliferative cells and tissues *in vivo*, and are proposed to arise from irreparable breaks or damage in telomeres ^9–12^.

The term TelOxidation was recently coined to describe the hypersensitivity of telomeres to oxidative stress ^13^. Oxidative stress arising from endogenous sources, including inflammation, and exogenous sources such as pollution, is not only associated with accelerated telomere shortening but also increased dysfunctional telomeres even in the absence of attrition ^14^. Telomeric TTAGGG repeats are preferred sites for the production of the highly common oxidative lesion 8-oxoguanine (8oxoG). Using a chemoptogenetic tool to selectively produce 8oxoG at telomeres, we demonstrated that the chronic and persistent formation of telomeric 8oxoG in HeLa LT cancer cells accelerates telomere shortening and drives hallmarks of telomere crisis including dicentric chromosomes ^15^. More recently, we showed that acute production of telomeric 8oxoG in non-diseased human fibroblasts and epithelial cells triggers rapid telomere dysfunction and p53-mediated cellular senescence in the absence of telomere shortening ^16^. We found 8oxoG induces telomere dysfunction by causing replication stress at telomeres, consistent with reports that 8oxoG can stall replicative polymerases ^17^. However, the role of 8oxoG processing in telomere stability and cellular outcomes remains incompletely understood.

Base excision repair (BER) acts to preserve the genome after 8oxoG formation ^18, 19^. OGG1 glycosylase specifically recognizes and removes 8oxoG opposite C, producing an apurinic/apyrimidinic (AP) site, which is primarily cleaved by AP endonuclease 1 (APE1), generating a nick and 5’-deoxyribose phosphate (5’dRP). DNA polymerase β (Pol β) removes the 5’dRP and fills the gap, then DNA ligase III seals the nick. 8oxoG is prone to mispairing with A if not repaired prior to DNA replication, causing G:C to T:A transversions ^20–23^. To prevent this, MUTYH glycosylase removes the undamaged A opposite 8oxoG, then polymerase lambda inserts a C to generate an 8oxoG:C basepair which is eventually restored to G:C by OGG1 mediated repair ^19, 24^. Poly (ADP-ribose) polymerases PARP1 or PARP2 bind AP and nick repair intermediates and synthesizes poly(ADP-ribose) PAR chains to recruit downstream BER proteins ^25^. XRCC1 interacts with Pol β and ligase 3, and prevents PARP1 trapping on DNA ^26^. Studies in mice for OGG1 and MUTYH, and in humans for MUTHY, established a role for unrepaired 8oxoG lesions in tumorigenesis ^27–29^. Furthermore, UV-DDB promotes turnover and dissociation of OGG1 and MUTYH, facilitating hand-off to APE1 and underscoring the importance of rapid and efficient repair progression ^30, 31^. While 8oxoG damage increases with age ^32^, potential roles for BER processing of telomeric 8oxoG in cellular aging are poorly understood.

Here we used our chemoptogenetic tool to selectively produce 8oxoG at telomeres in human fibroblasts singly or doubly deficient in OGG1 and MUTYH glycosylases. Surprisingly, we observed that knock out of OGG1 or MUTYH partially rescued multiple hallmarks of telomeric 8oxoG induced senescence, whereas double knock out caused a near complete rescue. Moreover, telomeric 8oxoG induced fewer fragile telomeres in single knock-out cells, compared to wild type, and virtually no increase in double knock-out cells. Consistent with the conversion of 8oxoG to single-strand breaks (SSB) intermediates, the prevention of BER initiation in OGG1 and MUTYH deficient cells suppressed telomeric 8oxoG induced PARylation and rendered cells insensitive to synergistic effects of PARP inhibitors on damage-induced senescence. However, OGG1 is required to protect cells from senescence caused by chronic telomeric 8oxoG, whereas MUTYH promotes senescence to prevent chromosomal instability from persistent damage over time. Collectively, our studies indicate that inefficient completion of BER at telomeres triggers cellular senescence via an accumulation of repair intermediates which impair telomere replication and stability.

## RESULTS

### OGG1 and MUTYH deficiency reduces sensitivity to acute oxidative telomere damage

To study how oxidative damage impacts telomere function, we use a chemoptogenetic tool that generates 8oxoG by directing localized production of highly reactive singlet oxygen (^1^O_2_) at telomeres ^15^. Photosensitizer di-iodinated malachite green (MG2I) dye (D) produces ^1^O_2_ when bound to a fluorogen activating peptide (FAP) fused with telomeric protein TRF1, and then activated with 660 nm light (L) ^33, 34^. We previously demonstrated that acute production of telomeric 8oxoG in non-diseased human BJ hTERT fibroblasts (BJ FAP-TRF1) increased senescent cells just 4 days after dye and light (DL) exposure ^16^. To determine the role of BER processing in modulating telomeric 8oxoG-induced senescence, we disrupted *OGG1* and *MUTYH* genes in BJ FAP-TRF1 cells. Western blotting confirmed OGG1 and MUTYH loss in single and double knock out (KO) cells, and similar FAP-TRF1 expression levels, indicating consistent targeted 8oxoG production among the cell lines (Figures 1A-B). Surprisingly, relative cell counts 4 days after 20 min of DL exposure showed a partial rescue of 8oxoG-induced growth reduction in single KO cells, and a near complete rescue in DKO cells, while all cell lines were unaffected by dye or light alone (Figure 1C). Increasing the DL exposure from 5 to 20 min amplified the differential response of glycosylase deficient and WT cells to telomeric damage (Figure S1A), consistent with greater 8oxoG production with longer exposures ^33^. Simultaneous treatment with 20 min DL and the OGG1 inhibitor TH5487 (OGG1i), which prevents OGG1 binding ^35^, also partially rescued the cell growth reduction in WT cells, similar to OGG1 KO, whereas a close structural analog (OGG1i^NA^) had no effect (Figure S1B). MUTYH KO cells treated with DL and OGG1i resembled the response of the DKO cells (Figure S1C). Since glycosylase deficiency alone reduced growth rates (Figure S1D), we asked if this contributed to the reduced sensitivity to telomeric 8oxoG. For this, we cultured WT cells in limiting serum to slow their growth. We observed a similar 8oxoG-induced decrease in relative cell number and no variation in the induction of cellular senescence (Figure S1E,F). These results confirm that the reduction in 8oxoG sensitivity is due to glycosylase deficiency and not to differences in growth rates among the cell lines.

**Figure 1.**
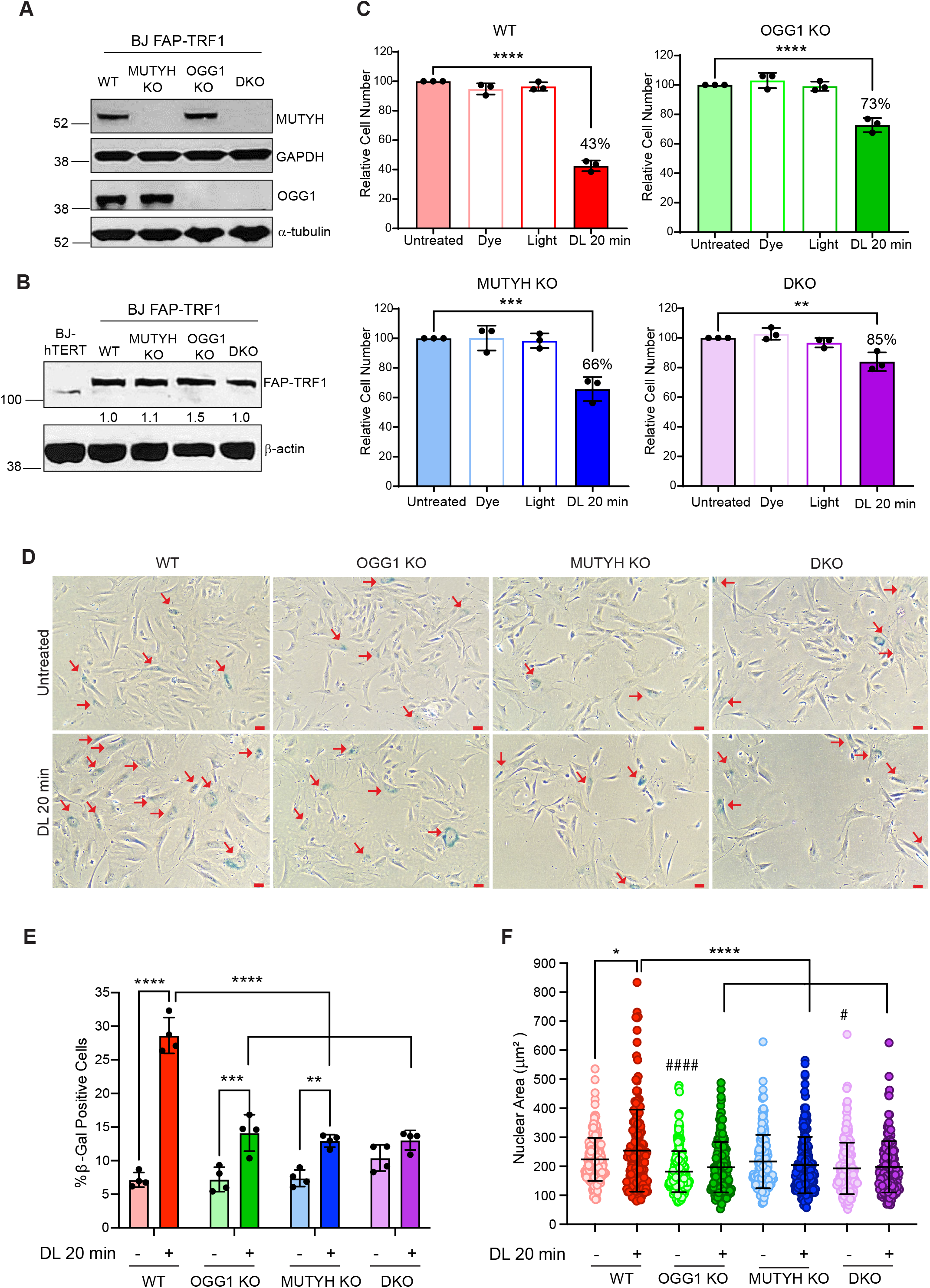
OGG1 and MUTYH deficiency reduces sensitivity to acute oxidative telomere damage. (A) Immunoblot of MUTYH and OGG1 in parental BJ FAP-TRF1 and MUTYH KO, OGG1 KO or DKO cell lines. α-tubulin and GAPDH used as loading controls. (B) Immunoblot of TRF1 in parental and BJ FAP-TRF1 cell lines. β-actin used as a loading control. (C) Cell counts of BJ FAP-TRF1 cell lines obtained 4 days after recovery from 20 min dye + light (DL) treatment, relative to untreated cells. Data are mean ± s.d. from 3 independent experiments; 2-way ANOVA; **p ≤ 0.01, ***p ≤ 0.001. ****p ≤ 0.0001. (D) Representative images of β-galactosidase positive cells obtained 4 days after recovery from no treatment or 20 min DL treatment. Red arrows mark positive cells. (E) Percent β-galactosidase positive cells. Data are mean ± s.d. from 4 independent experiments; 2-way ANOVA; **p ≤ 0.01; ***p ≤ 0.001; ****p ≤ 0.0001. (F) Size of nuclear area (μm^2^) of cells obtained 4 days after recovery from no treatment or 20 min DL. Data are mean ± s.d. from total nuclei analyzed; each dot represents a nucleus; ordinary one-way ANOVA; comparisons *p ≤ 0.05; ****p ≤ 0.0001; comparison of untreated cell lines with untreated WT indicated by #p ≤ 0.05; ###p ≤ 0.001.

Next, we asked whether the differential growth reduction after damage among the cell lines was due to differences in the induction of senescence ^36^. In contrast to WT cells, which showed a dramatic 4-fold increase in senescence-associated β-galactosidase (SA-β-gal) staining 4 days after 20 min DL exposure, the OGG1 and MUTYH KO cells showed a lesser (about 2-fold) increase, and the DKO cells remained largely unaffected (Figure 1D,E). Consistent with morphological changes of senescent cells ^37, 38^, the nuclear area of WT cells increased significantly 4 days after DL treatment, whereas the glycosylase deficient cells showed minimal changes (Figure 1F). Taken together, our data show that OGG1 or MUTYH deficiency in non-diseased cells reduces sensitivity to the acute telomeric 8oxoG induction of senescence-associated phenotypes.

### Glycosylase activity enhances telomeric 8oxoG-induced cytoplasmic DNA

Another hallmark of senescent cells is the appearance of cytoplasmic chromatin fragments (CCFs), which we previously showed increased in BJ FAP-TRF1 cells after telomeric 8oxoG damage ^16^. CCFs resemble micronuclei (MN) but arise independently of mitosis and chromosome breaks ^39^. As further confirmation that the repair deficient cells have an attenuated response to telomeric 8oxoG damage, the production of CCFs in the single KO and DKO cells was reduced or abrogated, respectively, 24 h and 4 days after DL (Figures 2A-B and Figure S2A). The content of telomeric DNA in the CCFs was comparable in the glycosylase proficient and deficient cells (Figure S2B), and the proportion of CCFs containing γH2AX was not increased by the targeted oxidative damage at telomeres in the single KO and DKO cells (Figure 2C). The percentage of CCFs colocalizing with 53BP1 was also not altered by dye and light treatment, and was significantly lower than the percent containing γH2AX for all the cell lines (Figure 2C). This is consistent with the cytoplasmic DNA species arising from chromatin blebbing rather than chromosome breakage as we reported previously for WT cells ^16^. In agreement, the oxidative telomere damage did not increase chromatin bridges in any of the cell lines (Figure 2D). This suggests that while the damage-induced cytoplasmic DNA species were decreased in glycosylase-deficient cells, the mechanism of production was the same as in WT cells. Since MN or CCFs can arise from DNA breaks related to apoptosis induction, we tested for apoptosis by Annexin V (AV) and propidium iodide (PI) staining. Our results confirmed that 20 min DL exposure did not induce apoptosis in any of the cell lines, unlike the UV-treated controls (Figure S2C,D). In summary, cells lacking OGG1 or MUTYH, or both, show reduced formation of telomeric 8oxoG induced senescence-associated cytoplasmic DNA species.

**Figure 2.**
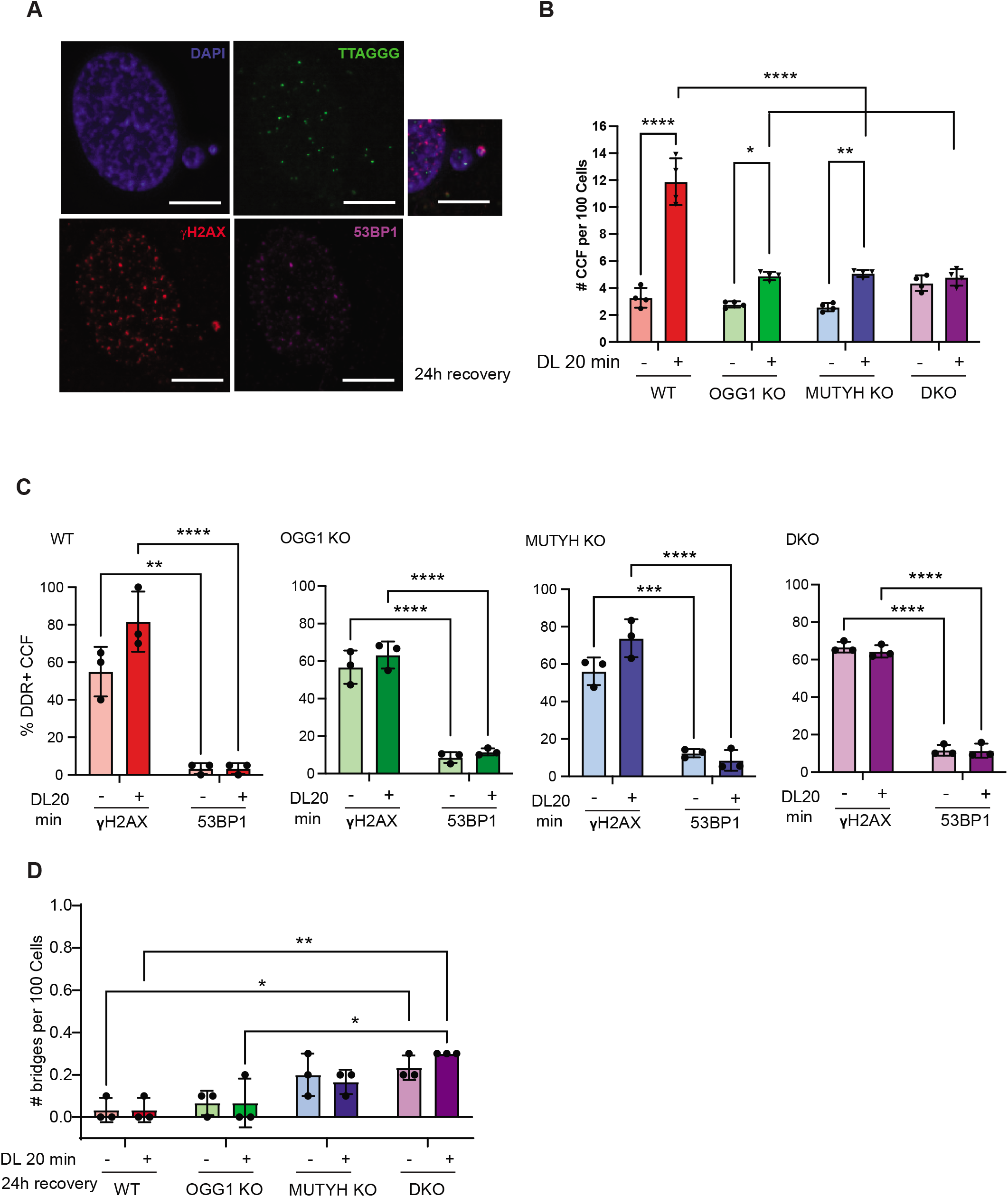
Glycosylase activity enhances telomeric 8oxoG-induced cytoplasmic DNA. (A) Representative IF-FISH image of cytoplasmic DNA species stained by DAPI (blue) containing telomeres (green), γH2AX (red) or 53BP1 (purple) in WT cells 24h after recovery from 20 min DL. (B) Quantification of cytoplasmic chromatin fragments (CCF) 24h after recovery from 20 min DL from panel A. Data are mean ± s.d. from 4 independent experiments, of at least 400 nuclei counted per experiment; 2-way ANOVA; *p ≤ 0.05, **p ≤ 0.01, ****p ≤ 0.0001. (C) Quantification of the percent of CCFs from panel (A) positive for DDR markers γH2AX or 53BP1 for the indicated cell lines. Data are mean ± s.d. from 3 independent experiments; 2-way ANOVA; **p ≤ 0.01, ****p ≤ 0.0001. (D) Quantification of the percent of chromatin bridges 24h after recovery from 20 min DL. Data are mean ± s.d. from 3 independent experiments, of at least 1,000 nuclei counted per experiment; 2-way ANOVA; *p ≤ 0.05, **p ≤ 0.01.

### Glycosylase deficiency suppresses telomeric 8oxoG induced replication stress

Telomeric 8oxoG induces senescence by impairing telomere replication, as indicated by the elevation of fragile telomeres and a localized DNA damage response (DDR) that ultimately activates p53-mediated premature senescence ^16^. We predicted that the greater senescence induction in WT cells compared to OGG1 and MUTYH deficient cells, was due to the glycosylases enhancing or provoking telomeric replication stress and subsequent DDR activation. To test this, we examined telomere integrity by fluorescent in situ hybridization (FISH) on metaphase chromosomes, and scored the chromatid ends as normal (one distinct telomeric foci), signal free (no staining) or fragile telomeres (multiple foci) (Figure 3A,B and Figure S3A). Transient depletion of p53 through siRNA (Figure S3B) allowed the damaged cells to progress into mitosis. While DL treatment did not significantly increase signal free ends in any of the cell lines (Figure 3A), telomeric 8oxoG induced fewer fragile telomeres in the single KO cells compared to WT, and virtually no increase in the DKO cells (Figure 3B). These results are consistent with the attenuated senescent phenotypes observed in the glycosylase deficient cells after damage, and suggest that OGG1 and MUTYH deficiency suppresses replication stress induced by telomeric 8oxoG.

**Figure 3.**
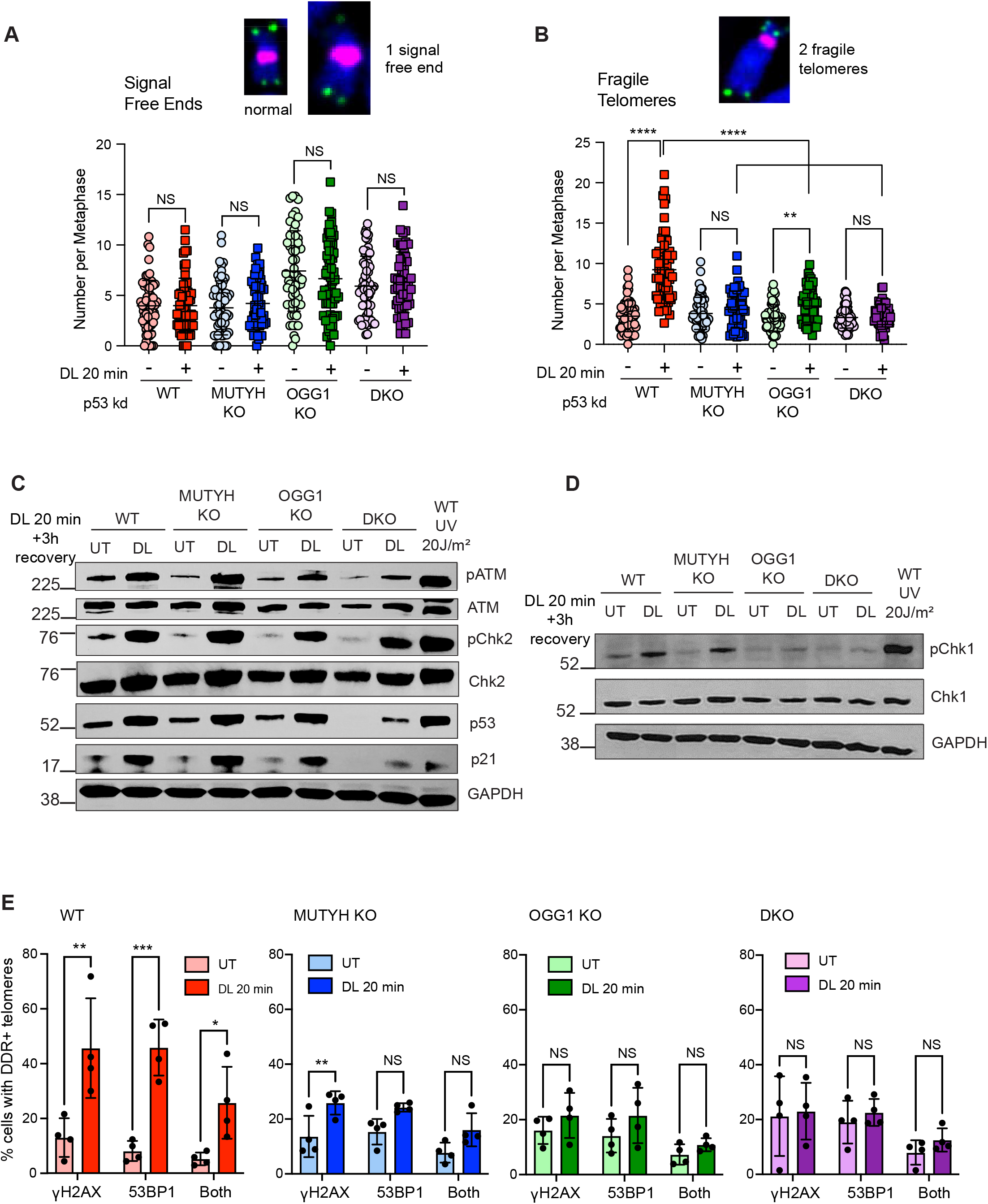
Glycosylase deficiency suppresses telomeric 8oxoG induced replication stress. (A-B) Number of telomeric signal-free chromatid ends (A) or fragile telomeres (B) per metaphase in cell lines transiently depleted for p53. Inset shows representative images of metaphase chromosomes scored as normal, signal free end or fragile by telo-FISH (green); pink = centromeres. Data represent mean ± s.d. from number of metaphases indicated as dots analyzed from three independent experiments, normalized to the chromosome number; two-way ANOVA; NS = not significant, **p ≤ 0.01, ****p ≤ 0.0001. (C) Immunoblot of indicated proteins in cells untreated or treated with 20 min DL, or 20 J/m^2^ UVC light as a positive control (WT only), and recovered 3h; pATM and pChk2 indicate phosphorylated forms; GAPDH used as a loading control. (D) Immunoblot of indicated proteins in cells untreated or treated with 20 min DL, or 20 J/m^2^ UVC light as a positive control (WT only), and recovered 3h; pChk1 indicates phosphorylated forms; GAPDH used as a loading control. (C) Quantification of the percent of cells showing ≥1 telomere foci co-localized with γH2AX, 53BP1 or both 24 h after no treatment or 20 min DL. Data represent the mean ± s.d. from four independent experiments, of more than 50 nuclei analyzed per condition for each experiment; 2-way ANOVA; NS = not significant (shown only for untreated to DL comparisons), *p ≤ 0.05; **p ≤ 0.01; ***p ≤ 0.001.

Telomeric replication stress and fragility lead to DDR signaling and activation of ATM and ATR kinases at the telomeres ^16, 40^. Therefore, we tested whether glycosylase deficiency also suppresses activation of the ATM/Chk2 and ATR/Chk1 pathways and downstream induction of p53 and p21, after telomeric 8oxoG damage. While both OGG1 and MUTYH KO cells showed ATM/Chk2, p53 and p21 activation shortly (3 h) after 20 min of DL, similar to WT cells, the DKO cells showed an attenuated response (Figure 3C). Phosphorylation of Chk1 after 8oxoG damage was attenuated in single knockout and DKO cells, compared to WT, although it was higher in MUTYH KO cells compared to the other glycosylase-deficient cells (Figure 3D). These results are consistent with less 8oxoG-induced replication stress in the glycosylase-deficient cells, compared to WT. As a positive control, we confirmed that all the cell lines mount a robust DDR after 20 J/m^2^ UVC exposure (Figure S3C). Thus, while lesion processing by either OGG1 or MUTYH can activate ATM or ATR signaling, the impaired DDR activation in the DKO cells suggests the BER initiation at telomeric 8oxoG lesions contributes to ATM/ATR kinase activation. The analysis of the localized telomeric DDR confirmed that 20 min DL treatment significantly increases recruitment of γH2AX and/or 53BP1 at telomeres 24 h after damage (Figure 3E and Figure S3D). However, telomeric 8oxoG damage induced only a slight increase in single KO cells showing DDR+ telomeres, and a lesser increase in DKO cells. In summary, the loss of both MUTYH and OGG1 glycosylases suppresses telomeric 8oxoG induction of DDR+ telomeres, DDR signaling, and p53 activation, consistent with the attenuated production of senescent cells.

### BER intermediates promote telomeric 8oxoG-induced senescence

Next, we examined the mechanism by which glycosylase deficiency attenuates the telomere damage-induced DDR and senescence. We hypothesized that the initiation of BER at telomeric 8oxoG lesions by OGG1 and MUTYH and the consequent formation of abasic sites and downstream SSBs may impair telomere replication and trigger cellular senescence. To test for the formation of SSB intermediates after telomeric 8oxoG production in the WT and glycosylase deficient cell lines, we examined Poly-ADP ribosylation (PARylation). PARP1 and PARP2 are recruited to SSBs and PARylate themselves and other repair proteins to promote BER completion ^25, 41^. PAR chains are degraded very rapidly by the Poly(ADP-Ribose) Glycohydrolase (PARG) ^42^. To preserve the PAR modification for detection, we treated cells during DL exposure with the PARG inhibitor PDD00017272 (PARGi). We observed weak, but detectable, PARylation after 5 min DL, and much more robust PARylation after 20 min DL (Figure 4A). Dye or light only controls failed to induce PARylation, confirming that PARP1/2 activity depends on the targeted 8oxoG formation at telomeres (Figure S4A). Remarkably, while PARylation in the glycosylase single KO cells was reduced after telomeric 8oxoG induction, compared to WT, it was nearly suppressed in DKO cells (Figure 4B). These data indicate the formation of BER intermediates in the DKO cells is negligible, and importantly, confirm that the DL treatment does not directly produce SSBs as the primary lesion. Furthermore, increased PARylation after 20 min DL, compared to 5 min, is consistent with the greater difference in growth inhibition between the WT and repair-deficient cell lines after 20 min damage compared to 5 min (compare Figure 1C and Figure S1A). The depletion of APE1 also partially rescued the reduction in cell growth and the increase in senescent cells caused by 20 min DL (Figures S4B-D). Since APE1 loss prevents the conversion of the abasic sites to an SSB, this suggests that SBB repair intermediates at telomeres may be more detrimental than abasic sites, leading to increased senescence.

**Figure 4.**
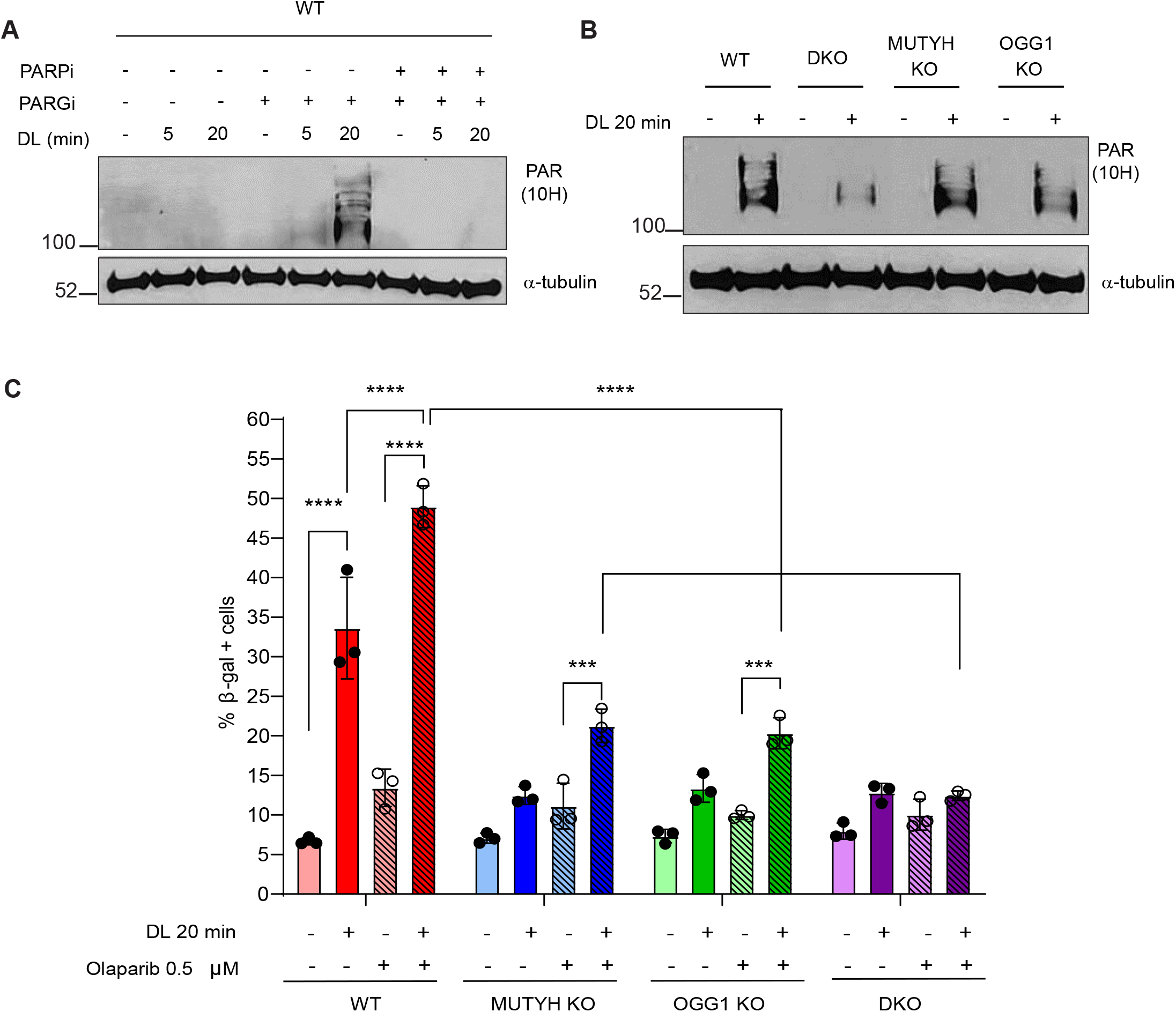
BER intermediates promote telomeric 8oxoG-induced senescence. (A) Immunoblot of poly-ADP ribose (PAR) in WT BJ FAP-TRF1 treated with 10 μM PARG inhibitor PDD 00017272 (PARGi), 10 μM PARP1 inhibitor AZD2281 (Olaparib - PARPi) and with DL for the indicated times. α-tubulin used as a loading control. (B) Immunoblot of PAR in cell lines treated with 10 μM PARGi and with 20 min DL. α-tubulin used as a loading control. (C) Percent β-galactosidase positive cells treated with 20 min DL with or without 0.5 μM Olaparib (PARPi). Data are mean ± s.d of 3 independent experiments; 2-way ANOVA; ***p ≤ 0.001, ****p ≤ 0.0001.

Next, we reasoned that if repair intermediates were present after 8oxoG damage, then the addition of the PARP inhibitor Olaparib should increase PARP1 or PARP2 retention at these sites, and further exacerbate replication stress and senescence induction ^43^. By enhancing PARP1/2 retention at SSBs, Olaparib causes aborted BER and accumulation of toxic repair intermediates ^43, 44^. Combined treatment of DL and Olaparib in BER proficient cells synergistically increased senescence induction compared to DL alone. Conversely, single knock out cells were less affected, and DKO cells were insensitive to adding the PARPi with damage (Figure 4C). Taken together, these results suggest that the initiation of BER and consequent formation of repair intermediates are drivers of telomeric 8oxoG-induced senescence.

### Both BER and replication stress activate PARylation and DDR after 8oxoG damage

Next, we investigated the mechanism of 8oxoG-induced PARylation and DDR activation. While SSB formation can activate both DDR and PARylation, we previously demonstrated that replication contributes to telomeric 8oxoG-induced DDR in WT cells ^16^, and other studies revealed PARylation of proteins at sites of replication stress ^45^. We synchronized the BJ fibroblasts to G0/G1 by serum starvation and contact inhibition to compare the response to telomeric 8oxoG in unsynchronized cells to quiescent (non-replicative) cells (Figure 5A). The absence of Cyclin A confirmed that cells seeded nearly at confluence and serum starved for 48 h enter quiescence (Figure S5A-B). We observed Chk2 phosphorylation in both unsynchronized and quiescent WT cells, indicating that telomeric 8oxoG processing by BER can trigger the DDR (Figure S5A). Telomeric 8oxoG damage induced robust PARylation in quiescent cells, although reduced compared to unsynchronized replicating cells (Figure 5B). Colocalization of PAR at telomeres in WT cells treated with PARGi revealed that telomeric 8oxoG generated PAR enrichment specifically at telomeres, which was attenuated in quiescent cells compared to unsynchronized cells (Figure 5C,D). These data indicate that replication stress induced by 8oxoG and its repair intermediates enhance PARylation, consistent with a role for PARPs in replication stress ^46^.

**Figure 5.**
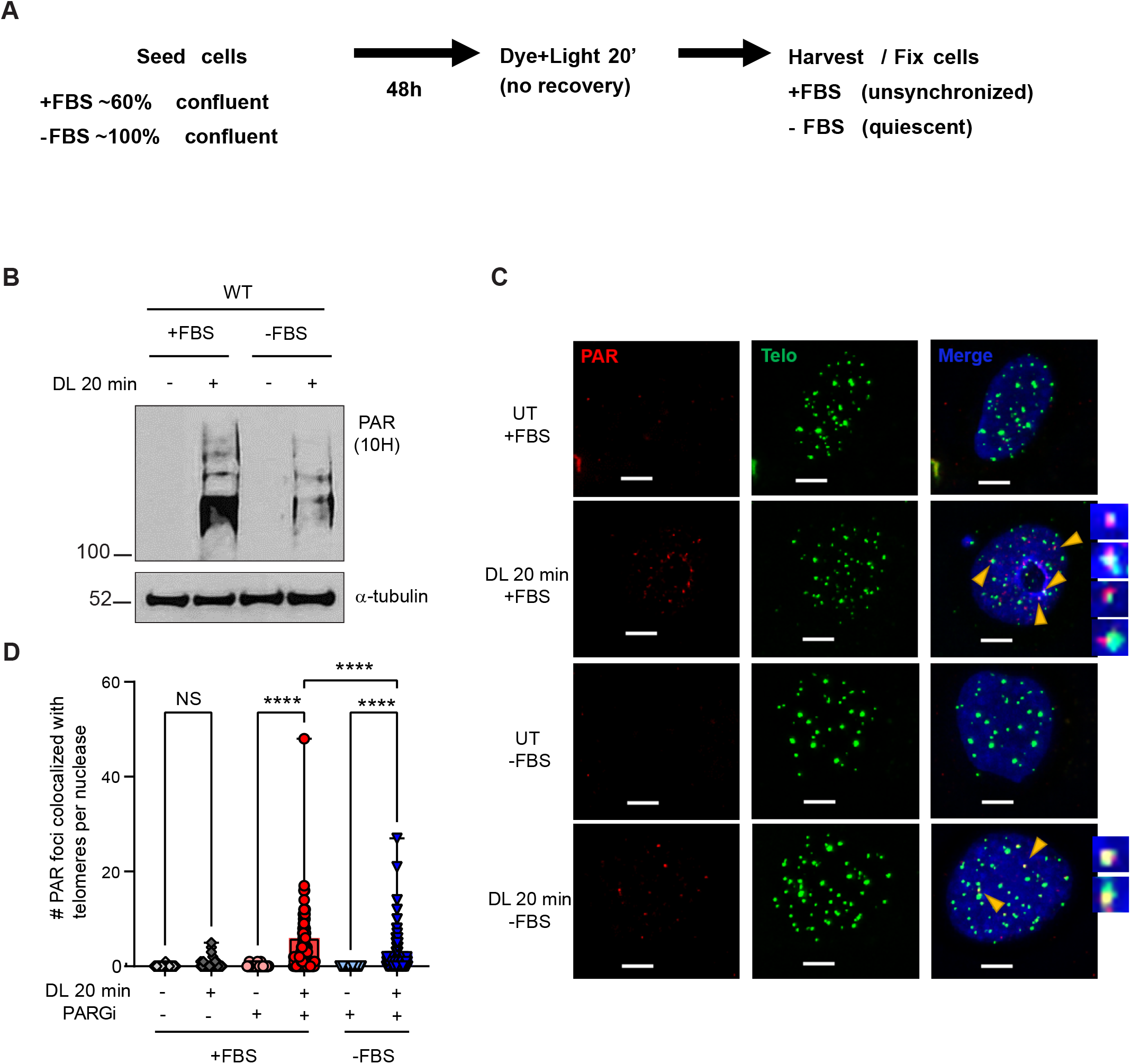
Both BER and replication stress activate PARylation after 8oxoG damage. (A) Schematic of experiments for PAR detection by WB (B) and telomere IF-FISH (C,D) in unsynchronized (cells grown with 10% FBS (+FBS)) and quiescent (cells grown with 0.1% FBS (–FBS)) WT cells. (B) Immunoblot of PAR unsynchronized (+FBS) or quiescent (-FBS) WT cells treated with 10 μM PARGi and with no treatment or after 20 min DL. α-tubulin used as loading control. (C) Representative IF-FISH images showing PAR (red) colocalizing with telomeres (green) by telo-FISH in + or –FBS WT cells with no treatment or after 20 min DL. Yellow arrowheads point to colocalization of PAR with telomeres. (D) Quantification of the number of PAR foci colocalizing with telomeres per nuclei analyzed; each dot represents a nucleus. Data are mean ± s.d; ordinary one-way ANOVA with Tukey’s multiple comparisons test; NS = not significant (show only for untreated and DL comparisons); ****p < 0.0001.

### OGG1 loss sensitizes cells to chronic telomere damage while MUTYH loss promotes resistance

Here we show OGG1 or MUTYH glycosylase deficiency can partially suppress the deleterious effects of acute oxidative telomere damage in fibroblasts. However, we found previously that OGG1 repair of telomeric 8oxoG is critical for telomere stability in cancer cells after chronic telomere damage ^15^. Therefore, we investigated how repeated exposure to targeted 8oxoG impacted telomere stability and cellular growth in the BJ FAP-TRF1 fibroblasts. To mimic chronic oxidative stress at telomeres, we treated WT and p53 KO cells with DL each day for 10 min, except every fourth day when cells were harvested for analysis, over 24 days for 18 treatments (N18). Repeated damage dramatically decreased population doubling (PD) and increased senescence of the WT cells, while p53 KO cells were less affected, consistent with p53-mediated senescence and a similar lesser growth reduction in p53 negative HeLa cancer cells ^15^ (Figure S6A-C). Next, we examined how glycosylase deficiency impacted the response to repeated telomeric 8oxoG induction over 24 days (N18) with 20 min DL. Interestingly, loss of MUTYH or OGG1 alone reduced growth compared to WT cells, but the DKO cells grew nearly as well as WT (Figures 6A,B). Repeated DL exposure reduced growth of all the cell lines. However, the WT and OGG1 KO cells were the most severely affected as their growth curves began to plateau near day 12 or 8, respectively, while the MUTYH KO cells showed reduced but sustained growth over time (Figures 6A-C). Although the DKO cells also showed a significant PD decrease in response to the chronic telomeric damage, they surprisingly continued growing without reaching a plateau (Figure 6A). Between days 20 and 24 of damage, the growth curve for the DKO cells surpassed the WT cells. Consistent with the growth results, chronic telomeric 8oxoG damage induced a significant increase in β-gal positivity in WT and OGG1 KO cells, which was attenuated in the MUTYH KO and DKO cells (Figure 6D). This suggests the loss of MUTYH, but not OGG1, partially rescues senescence caused by chronic oxidative telomere damage, leading to reduced but sustained growth.

**Figure 6.**
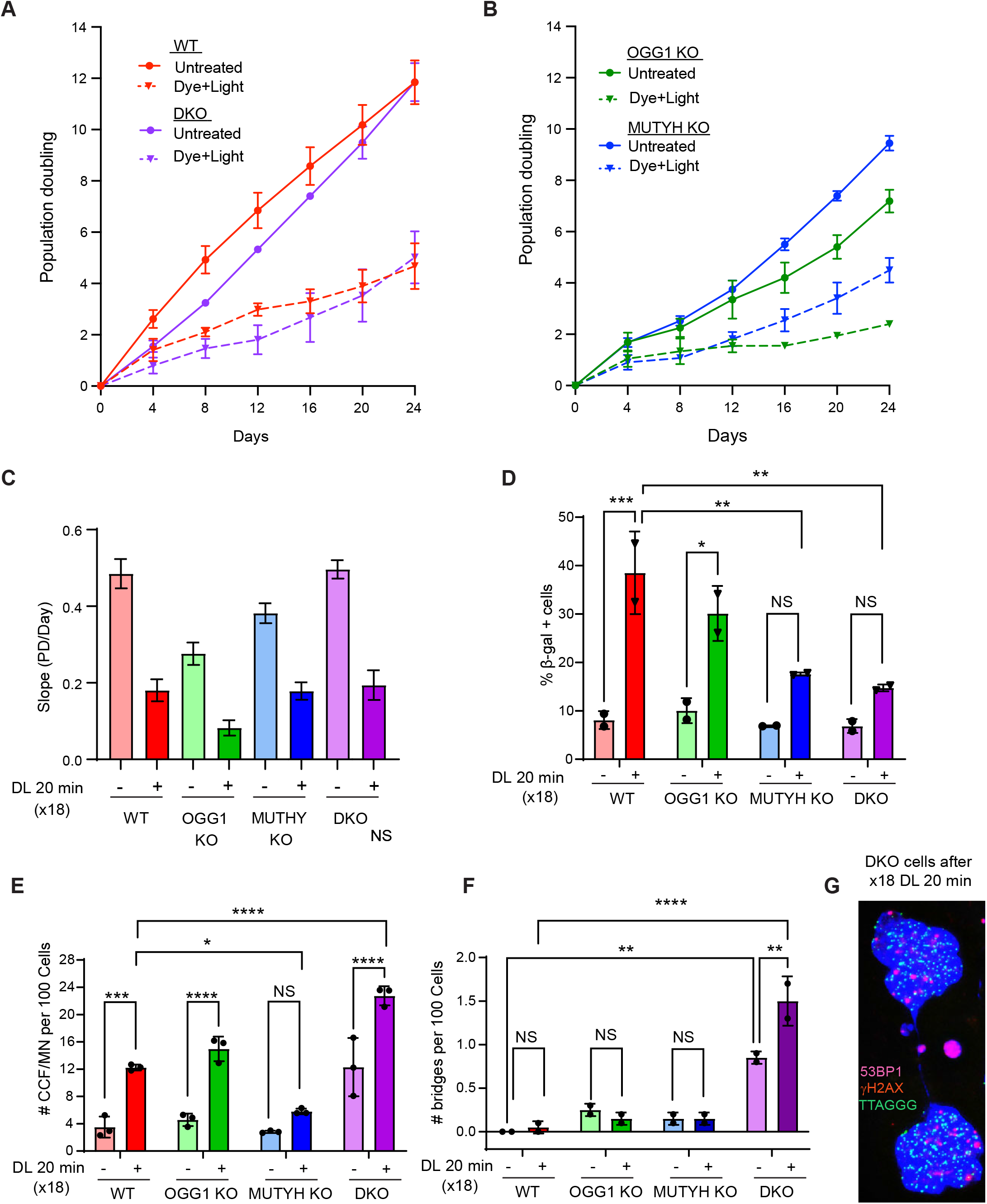
OGG1 loss sensitizes cells to chronic telomere damage while MUTYH loss promotes resistance. (A-B) Population doubling (PD) over 24 days of untreated cells (solid line) and cells treated with DL for 20 min each day except every 4^th^ day of harvest (dotted line). Data are mean ± s.d. from 3 independent experiments. (C). Simple linear regression showing the slope (PD/Day) of cells treated 18 times with 20 min DL. Error bars represent the 95% confidence intervals. (D) Percent β-galactosidase positive cells exposed for 18 times to 20 min DL over 24 days. Data are mean ± s.d. from 2 independent experiments; 2-way ANOVA; NS = not significant (shown only for untreated to DL comparisons);* p ≤ 0.05, **p ≤ 0.01, ***P ≤ 0.001. (E) Quantification of CCF/MN foci 24 h after recovery from last exposure to 20 min DL (x18). Data are mean ± s.d. from 3 independent experiments, of at least 400 nuclei scored per each experiment; 2-way ANOVA; NS = not significant (show only for untreated to DL comparisons), *p ≤ 0.05, ***p ≤ 0.001, ****p ≤ 0.0001. (F) Quantification of the number of bridges per 100 DKO cells counted. Data are mean ± s.d. from 2 independent experiments, of at least 1,000 nuclei counted per experiment; 2-way ANOVA; NS = not significant (shown only for untreated to DL comparisons); **p ≤ 0.01, ****P ≤ 0.0001. (G) Representative image of micronuclei and chromatin bridge in DKO cells after 24h from last of 18 exposures.

Next, we examined telomere integrity and cytoplasmic DNA production after chronic oxidative telomere damage. We observed a significant induction of cells with DDR+ telomeres in the WT and OGG1 KO cells 24 h after the last DL treatment (N18) (Figures S6D-G). Likewise, 18 DL exposures triggered a significant increase in cytoplasmic DNA species in WT and OGG1 KO, but not in MUTYH KO cells (Figure 6E). Interestingly, while the DKO cells showed less β-gal staining, they showed the largest increase in cytoplasmic DNA after damage, and the highest basal level prior to treatment (Figure 6E). This contrasts the failure of acute telomeric 8oxoG to increase cytoplasmic DNA species in the DKO cells (Figure 2B). Futhermore, the DKO cells also exhibited an increase in chromatin bridges 24h after N18, which was not detected in the WT or single KO cells, or after acute damage (Figures 6F,G). This suggests that some of the induced cytoplasmic DNA species in DKO cells may have arisen from chromatin bridge formation and breakage rather than senescence-associated blebbing from the nucleus. While telomeric 8oxoG did not alter the percent of total cyctoplasmic DNA species that stained positive for γH2AX or 53BP1 (Figure S6H), the DKO cells showed a higher percent with 53BP1, compared to WT or single knock-out cells. Taken together, these data suggest that glycosylase proficient fibroblasts undergoing telomeric oxidative stress experience enforcement of cellular senescence overtime. While OGG1 deficiency partially protects against acute telomeric 8oxoG, it becomes detrimental after long term damage. Moreover, MUTYH deficiency promotes resistance to chronic telomeric 8oxoG, while rescuing senescence induction and further promoting genome instability in cells simultaneously deficient for OGG1.

## DISCUSSION

Telomere oxidative damage has a direct role in cellular aging, as a single induction of telomeric 8oxoG is sufficient to induce hallmarks of cellular senescence in non-diseased cells ^16^. Here, we demonstrate that OGG1 and MUTYH glycosylase activity triggers senescence via an accumulation of repair intermediates, which impair telomere replication and stability. Using our chemoptogenetic tool to selectively generate 8oxoG lesions at telomeres in human fibroblasts singly or doubly deficient in OGG1 and MUTYH glycosylases, we observed a partial or near complete rescue, respectively, of multiple hallmarks of telomeric 8oxoG-induced senescence. This is consistent with a mechanistic model in which the production of SSB intermediates during repair contributes to replication stress and DDR activation at telomeres, leading to p53-mediated senescence. In support of this model, we uncovered a synergistic effect of telomeric 8oxoG damage and PARP inhibition on inducing cellular senescence, to which glycosylase deficient cells are insensitive. However, the chronic production of telomeric 8oxoG revealed divergent roles for OGG1 and MUTYH in suppressing or promoting, respectively, damage-induced senescence.

Our findings on 8oxoG processing are consistent with previous work on alkylation lesions that demonstrated excessive glycosylase activity and unbalanced BER activity can be detrimental. Preventing BER initiation in mice deficient for alkyladenine DNA glycosylase (Aag^-^/^-^) suppresses methyl methanesulfonate (MMS)-induced cerebellar toxicity, while Aag overexpression exacerbates, and PARP1 loss partially abrogates, these phenotypes ^47, 48^. Increased PARP1 activity from an accumulation of BER intermediates can cause cell death via depletion of the critical metabolite nicotinamide adenine dinucleotide (NAD^+^) ^49, 50^, which has also been implicated in cellular aging and senescence ^51^. Moreover, excessive PARP1 retention renders repair intermediates inaccessible to downstream BER enzymes and increases sensitivity to alkylation base damage in XRCC1^−^/^−^ cells, since XRCC1 prevents PARP1 trapping ^26^. Previous work was limited to alkylation damage due to the inability to induce 8oxoG in the absence of general oxidant treatment. Overcoming this obstacle with our chemoptogenetic tool, we now show that 8oxoG production exclusively at the telomeres is sufficient to provoke robust PARylation, and that loss of both OGG1 or MUTYH is required to suppress SSB formation and PARylation. Our finding that the PARP inhibitor Olaparib exacerbates the telomeric 8oxoG-induced cellular senescence in cells able to initiate BER, indicates that processing telomeric 8oxoG lesions contributes to senescence induction. This work reveals that the detrimental effects of BER intermediates arising from processing alkylation base damage, now also apply to those arising at the common 8oxoG lesion. Our findings raise the possibility that the combined use of conventional chemotherapeutic agents, which cause oxidative stress in normal tissues ^52^, and PARP inhibitors, may contribute to premature cellular senescence and aging in cancer patients through a telomere-mediated mechanism.

Previous studies reported that enzymatic activity from N-methylpurine-DNA glycosylase (MPG) and uracil DNA glycosylase 2 (UNG2) activate ATM/Chk2 by cooperating with APE1 endonuclease to convert glycosylase-specific lesions into SSB intermediates ^53, 54^. MPG and APE1 activity following MMS treatment led to PARylation and ATM activation, in both replicating and quiescent cells, suggesting DDR activation can arise directly from SSB formation or as a consequence of SSB conversion to DSBs during replication ^54^. In these studies, the use of H_2_O_2_ to induce oxidative lesions was confounded by findings that oxidative stress can also activate ATM via direct protein oxidation ^55, 56^. In our study, we show that in absence of general oxidative stress, the processing of the oxidative lesion 8oxoG at telomeres can activate the ATM/Chk2 pathway even in non-replicating cells. This DDR activation might arise due to BER initiation caused by glycosylase excision, or by the downstream formation of repair intermediates and recruitment of PARP1. Interestingly, since Gs only exist on the telomere lagging strand, excision should only occur on one strand in non-replicating cells by OGG1, ruling out the possibility that excision on opposite strands produces a DSB that activated ATM ^57^. By suppressing the possibility of DSB formation from replication fork collision with an SSB, or from excision of clustered bases on opposing strands, our data strengthen the evidence that SSBs can directly activate ATM kinase and DDR.

Among the most surprising results of this study are the similar partial rescue of the induced senescence phenotypes when either OGG1 or MUTYH is depleted, and the requirement to deplete both glycosylases for near-complete rescue. MUTYH excises A misinerted opposite 8oxoG to prevent the accumulation of G:C→T:A transversion mutations that would be fixed in the genome after another replication cycle ^58^. Thus, we predicted that MUTYH knock out would cause a weaker rescue of cellular senescence after acute telomeric 8oxoG damage compared to OGG1 deficiency. However, we observed a similar rescue of all the analyzed phenotypes, and an additive effect when both glycosylases were knocked out. Interestingly, the mutation G396D located in the MUTYH 8oxoG recognition domain is among the most prevalent in MUTYH-associated polyposis (MAP) cancers ^58^, which suggests that the recognition of 8oxoG itself is a crucial MUTYH property. Consistent with our finding of telomeric 8oxoG-induced PARylation in OGG1 ko cells, others reported rapid MUTYH-mediated PARylation after oxidative damage, and increased 8oxoG levels in MUTYH deficient cells shortly after oxidative damage ^59, 60^. Thus, a role for MUTYH response to oxidative damage even prior to significant 8oxoG:A mispairing has been proposed to arise from MUTYH binding unproductively to 8oxoG and thereby impacting 8oxoG:C lesion processing and possibly replication fork progression ^28^. The potential for MUTYH to bind unproductively to 8oxoG:C may explain why MUTYH deficient cells show a growth advantage similar to OGG1 deficient cells after oxidative telomere damage.

While loss of glycosylase activity provided a growth advantage after acute telomere damage, the absence of OGG1 during the long-term telomeric 8oxoG production was highly detrimental, whereas the loss of MUTYH was not. We previously demonstrated that persistent telomeric 8oxoG damage in HeLa cells caused telomere shortening and loss, and impaired cell growth, which was greatly exacerbated by OGG1 deficiency ^15^. Here we found that non-diseased human fibroblasts are much more sensitive to chronic telomeric 8oxoG, primarily due to p53 activation, since fibroblasts lacking p53 experienced a moderate damage-induced growth reduction similar to HeLa cells. Remarkably, the additional loss of MUTYH could partially rescue the severe senescence induction and growth impairment experienced in OGG1 deficient cells after chronic telomere damage. One possibility is that unrepaired 8oxoGs that accumulate at telomeres in the absence of OGG1 could lead to both 8oxoG:C and 8oxoG:A basepairs depending on which nucleotide is inserted during replication ^61^. MUTYH may impair replication both by generating SSBs at 8oxoG:A basepairs and by binding non-productively to 8oxoG:C basepairs. The loss of MUTYH is known to promote tumorigenesis by increased mutations as well as by avoidance of cell death under oxidative stress ^62^. Here we showed that MUTYH loss partially suppresses senescence after both acute and chronic telomeric oxidative damage, and very interestingly, greatly increases genomic instability in cells lacking OGG1. This suggests that normal cells with mutated MUTYH exposed to oxidative stress may become cancerous not only from accumulation of mutations, but also from telomere-mediated senescence evasion and cellular transformation over time due to genomic instability.

Given that senescence is associated with pro-inflammatory responses, our findings have important implications for the role of OGG1 inhibitors in suppressing inflammation by impairing OGG1 recruitment to regulatory regions of proinflammatory cytokines in TNFα-challenged cells ^35^. OGG1 inhibitors also mitigate bleomycin-iduced pulmonary fibrosis and bacterial lung infections in mice, by regulating and reducing pulmonary inflammatory responses ^63, 64^. Our data suggest that OGG1 inhibitors may also suppress inflammation by preventing senescence caused by oxidative stress at telomeres. We found that inhibiting OGG1 mitigates senescence caused by acute telomeric 8oxoG damage, and therefore, should reduce the senescence-associated secretory phenotype (SASP) we observed previously in repair proficient cells ^16^. However, our finding that OGG1 loss greatly exacerbates senescence caused by chronic telomeric 8oxoG damage, indicates that anti-inflammatory therapies based on OGG1 inhibition could be unsuitable for chronic inflammatory conditions.

In summary, our study demonstrates that repair intermediates arising at telomeres from OGG1 and MUTYH glycosylase activity impair telomere replication and activate both PARylation and DDR at telomeres, contributing to 8oxoG-mediated cellular senescence.

## ACKNOWLEDGEMENTS

We are grateful to Brigitte Schmidt and Bruce Armitage (Carnegie Mellon University) for providing MG2I dye, and to the late Marcel Bruchez, to whom this manuscript is dedicated, for his exceptional collaboration and development of the chemoptogenetic tool. We thank Elise Fouquerel (University of Pittsburgh) for critical reading of the manuscript, and Prasanth Nyalapatla (University of Pittsburgh) for synthesis of the active and inative OGG1 inhibitor. We thank Fatimah Adisa and Madalyn Fry for technical assistance with some western blotting experiments. This work was supported by NIH grants F32AG067710-01, K99ES033771 (to R.P.B.), R01NS117000 (to P.W.), and R35ES030396 and R01CA207342 (to P.L.O). This project used the UPMC Hillman Cancer Center Cytometry Facility which is supported in part by award P30CA047904.

## AUTHOR CONTRIBUTIONS

M.D.R. and P.L.O conceived the study and designed the experiments. M.D.R. performed most of the experiments, except R.P.B conducted the chronic experiment comparing WT and p53 knock out cells and generated the BJ FAP-TRF1 WT, p53 knock out, and OGG1 knock out cells. P.W. and P.R.N. synthesized the active and inactive OGG1 inhibitors. M.D.R. and P.L.O. wrote the manuscript with assistance from the other authors.

## DECLARATION OF INTERESTS

The authors have no competing interests to declare.

## SUPPLEMENTARY INFORMATION

### Legends

**Figure S1. OGG1 and MUTYH deficiency reduces sensitivity to acute oxidative telomere damage. Related to Figure 1.**

(A) Cell counts of BJ FAP-TRF1 cell lines obtained 4 days after recovery from 5 min DL treatment, relative to untreated cells. Data are mean ± s.d. from 3 independent experiments; 2-way ANOVA; ***p ≤ 0.001, ****p ≤ 0.0001.

(B-C) Cell counts of WT (B) and MUTYH KO (C) obtained 4 days after recovery from 20 min DL treatment in presence of DMSO or 20 μM of OGG1 inhibitor TH5487 (OGG1i) or 20 μM of inactive OGG1 inhibitor (OGG1i^NA^), relative to cells not treated with DL. Data are mean ± s.d. from 3 independent experiments; 2-way ANOVA; *p ≤ 0.05, **p ≤ 0.01, ***p ≤ 0.001, ****p ≤ 0.0001.

(D) Cell counts of untreated cell lines obtained after 4 days of growth, normalized to WT cells. Data are mean ± s.d. from 3 independent experiments; ordinary one-way ANOVA with Tukey’s multiple comparisons test; NS = not significant, *p ≤ 0.05.

(E) Cell counts of WT cells cultured in the indicated FBS concentrations, obtained 4 days after recovery from 20 min DL treatment, relative to WT untreated cells grown in 10% FBS (basal conditions). Data are mean ± s.d. from 4 independent experiments; 2-way ANOVA; ***p ≤ 0.001, ****p ≤ 0.0001.

(F) Percent β-galactosidase positive WT cells cultured in the indicated FBS concentrations. Data are mean ± s.d. from 4 independent experiments; 2-way ANOVA; ****p ≤ 0.0001.

**Figure S2. Glycosylase activity enhances telomeric 8oxoG-induced cytoplasmic DNA. Related to Figure 2.**

(A) Quantification of cytoplasmic DNA species (CCFs) 4 days after recovery from 20 DL. Data are mean ± s.d. from 3 independent experiments, of at least 400 nuclei scored per experiment; 2-way ANOVA; *p ≤ 0.05, ****p ≤ 0.0001.

(B) Quantification of CCFs positive for telomeric DNA staining 24 h after recovery from 20 min DL.

(C) Percent of cells positive for annexin V (AV), propidium iodide (PI), or both, 4 days after indicated treatments. Data are the mean ± s.d. from three independent experiments; two-way ANOVA; ****p < 0.0001.

(D) Representative scatterplots of Annexin V (x-axis) and propidium iodine (y-axis) staining of cells 4 days after the indicated treatments.

**Figure S3. Glycosylase deficiency suppresses telomeric 8oxoG-induced replication stress. Related to Figure 3.**

(A) Representative images of telomere FISH on metaphase spreads in p53 kd BJ FAP-TRF1 WT cells untreated or treated with 20 min DL. Green foci are telomeres and pink foci are CENPB centromeres. Scale bars = 10 μm. Orange and blue arrowheads point to signal free ends and fragile telomeres, respectively.

(B) Immunoblot of p53 in WT cells transiently transfected with siRNAs against p53 and cultured for the indicated times. GAPDH used as a loading control.

(C) Immunoblot of pChk1 and Chk1 in WT and MUTYH KO, OGG1 KO or DKO BJ FAP-TRF1 cells untreated or treated with 20 min DL, or 20 J/m^2^ UVC light as a positive control (WT only), and recovered 3 h. GAPDH used as a loading control.

(D) Representative IF images showing γH2AX (red) and 53BP1 (purple) staining with telomeres (green) by telo-FISH for WT cells 24 h after no treatment or 20 min DL. Yellow arrowheads point to NIS-Elements-defined intersections between 53BP1 and/or γH2AX with telomeres. Scale bars, 10 μm.

**Figure S4. BER intermediates promote telomeric 8oxoG-induced senescence. Related to Figure 4.**

(A) Immunoblot of PAR in WT BJ FAP-TRF1 cells treated with 10 μM PARGi and dye only, 20 min of light only or 20 min DL. α-tubulin used as a loading control.

(B) Immunoblot of OGG1 (top panel) and APE1 (bottom panel) at the indicated times following treatment of WT BJ FAP-TRF1 cells with targeting siRNA against OGG1 and APE1, respectively. α-tubulin used as a loading control.

(C-D) Cells were treated with targeting siRNA to knock down (kd) OGG1 or APE1, and then 24 h later treated with 20 min DL. (C) cell counts of WT cell lines obtained after 4 days after recovery from 20 min DL treatment, relative to untreated cells. Data are mean ± s.d. from 3 independent experiments; 2-way ANOVA; ***p ≤ 0.001. ****p ≤ 0.0001. (D) Percent β- galactosidase positive cells 4 days after recovery from DL20 min. Data are mean ± s.d. from 4 independent experiments; 2-way ANOVA; **p ≤ 0.01; ***p ≤ 0.001; ****p ≤ 0.0001.

**Figure S5. Both BER and replication stress activate PARylation after 8oxoG damage. Related to Figure 5.**

(A) Immunoblot of Cyclin A, phosphorylated Chk2 (pChk2) and Chk2 in WT cells untreated or treated with 20 min DL or 20 J/m^2^ UVC light as a positive control, and recovered 3 h. GAPDH used as a loading control.

(B) Quantification of EdU-positive cells in WT cells 3 hours after 20 min DL in 10% FBS (unsynchronized) or 0.1% FBS (quiescent). Error bars represent the mean ± s.d. from number of independent images analyzed.

**Figure S6. OGG1 loss sensitizes cells to chronic telomere damage while MUTYH loss promotes resistance. Related to Figure 6.**

(A) Population doubling (PD) over 24 days of untreated and repeatedly DL 10 min-treated WT and p53 KO cells. Data are means ± s.d from 6 (WT) and 3 (p53 KO) independent experiments.

(B) Simple linear regression showing the slope (PD/Day) of WT and p53 KO cells treated 18 times with DL 10 min. Error bars represent the 95% confidence intervals.

(C) Percent β-galactosidase positive WT and p53 KO cells exposed for 9 or 18 times to 10 min DL. Data are the mean ± s.d. from 3 (WT) and 2 (p53 KO) independent experiments.

(D-G) Quantification of the percentage of cells exhibiting telomere foci co-localized with γH2AX, 53BP1 or both for the indicated BJ FAP-TRF1 cell lines 24 h after N18 treatment of 20 min DL exposures. Data are the mean ± s.d. from three independent experiments, of more than 50 nuclei analyzed per condition for each experiment; two-way ANOVA; NS = not significant (shown only for untreated to DL comparisons), *p < 0.05.

(H) Quantification of the percent of cytoplasmic DNA species CCF/MN positive for DDR markers γH2AX or 53BP1 for the indicated cell lines. Data are mean ± s.d. from 2 independent experiments; 2-way ANOVA; **p ≤ 0.01, ****p ≤ 0.0001.

## METHODS

### Cell Culture and Cell Lines

BJ hTERT FAP-mCER-TRF1 wild type (BJ FAP-TRF1 WT) were previously described ^16^ and were grown in DMEM (Gibco) with 10% Hyclone FBS, 1% penicillin/streptomycin and 500µg/ml G418 (to maintain FAP-TRF1 expression). To generate knockout cell lines, HEK 293T cells were transfected with pLentiCRISPR V2 vectors (GenScript) expressing S. pyogenes Cas9 and guide RNAs designed and validated for uniquely targeting the human OGG1 and MUTYH genes ^65^, and with Mission Packaging Mix (Sigma) to produce lentivirus. Recipient cells were infected with virus collected 48 and 72 h post transfection and then selected as described below. OGG1 knock out (KO) and MUTYH KO cells were obtained by infection of BJ FAP-TRF1 WT with lentivirus expressing respectively guide RNAs targeting OGG1 exon 4 (gRNA3, sequence GCTACGAGAGTCCTCATATG) and MUTYH exon 2 (gRNA5, sequence GCATGCTAAGAACAACAGTC) and selected with 1.5 µg/ml Puromycin (Gibco). OGG1 KO/MUTYH KO (DKO) cells were obtained by infection of BJ FAP-TRF1 OGG1 KO cells with lentivirus expressing same MUTYH targeting guide RNA as described above, and were selected with 10 µg/ml Blasticidin S HCl (Gibco). After selection and death of uninfected cells, the infected cells were expanded and tested for mycoplasma, and expression of targeted protein(s) was determined by western blotting. Except for HEK 293T cells, all cells are maintained at 5% O_2_.

### Cell Treatments

To generate singlet oxygen at telomeres, cells were treated with dye and light (DL). Briefly, cells were plated at an appropriate density for the experiment overnight. The next day, cells were incubated in OptiMEM (Gibco) at 37 ° C for 15 min before adding 100nM MG2I for another 15 min. Cells were then exposed in the lightbox to a high intensity 660 nm LED light at 100 mW/cm^2^ for 20 min (unless indicated otherwise) to trigger excitation of the FAP-bound MG2I dye and the production of singlet oxygen. For chronic exposure to telomeric singlet oxygen, cells were treated with 10 or 20 min DL as described above, for three consecutive days and harvested/reseeded every fourth day for a total of 24 days and 18 exposures. PDD00017272 (PARGi) was added in OptiMEM at the indicated concentrations for the entire time of incubation with OptiMEM and DL treatment. AZD2281 (olaparib) was added in complete media at the indicated concentrations after DL treatment and cells were incubated for 4 days.

### Growth Analyses

For cell counting experiments, 25,000 cells were plated in 6-well overnight. Cells were treated as indicated and returned to the incubator and recovered for 4 days. Cells were detached from the plates, resuspended, and counted on a Beckman Coulter Counter. Each experiment had 3 technical replicates, which were averaged.

### Senescence Associated Beta-Galactosidase Assay

Detection of β-gal activity was done according to the manufacturer’s instructions (Cell Signaling). Briefly, cells were washed with PBS, and then fixed at room temperature for 10 minutes. Cells were washed again twice with PBS and then were incubated overnight at 37 °C with X-gal staining solution with no CO_2_. Images were acquired with a Nikon brightfield microscope with DS-Fi3 camera and analyzed in NIS Elements. At least 300-800 cells were counted per condition for each experiment.

### Immunofluorescence and FISH

Cells were seeded on coverslips and treated as indicated. Following treatment and/or recovery, cells were washed twice with PBS and fixed at room temperature with 4% formaldehyde for 10 min, except for PAR (10H) staining fixation was on ice in -20 °C methanol/acetone (v/v) for 30 min. Fixed cells were rinsed with 1% BSA in PBS, and washed 3x with PBS-Triton 0.2% before blocking with 10% normal goat serum, 1% BSA, and 0.1% Triton-x. Cells were incubated overnight at 4 °C with indicated primary antibodies. Next day cells were washed with PBS-T 3x before incubating with secondary antibodies and washing again 3x with PBS-T. To detect EdU incorporation, both unsynchronized and quiescent cells seeded on coverslips were pulsed with 20 μM EdU after 2h from 20 min DL treatment and incubated for an additional hour before being fixed and then Click chemistry was performed according to the manufacturer’s instructions (Thermo). When performing FISH, cells were re-fixed with 4% formaldehyde and rinsed with 1% BSA in PBS, and then dehydrated with 70%, 90%, and 100% ethanol for 5 min. Telomeric PNA probe was diluted 1:100 (PNA Bio, F1004, TelC-Alexa 488, 3xCCCTAA) in hybridization buffer (70% formamide, 10mM Tris HCl pH 7.5, 1x Maleic Acid buffer, 1x MgCl_2_ buffer) and boiled for 5 min at 85°C before returning to ice. Coverslips were incubated for 10 min on a hot plate at 75°C and then at room temperature for 2 h in humid chambers in the dark. After two washes in hybridization wash buffer (70% formamide, 10 mM tris HCl pH 7.5), coverslips were rinsed in water before staining with DAPI and mounting with ProLong™ Diamond Antifade Mountant (Thermo Fisher). Image acquisition was performed with a Nikon Ti inverted fluorescence microscope. Z stacks of 0.2 μm (60x objective) or 0.5 μm (20x objective) thickness were captured and images were deconvolved using NIS Elements Advance Research software algorithm. For MN counting at least 400 nuclei were scored per experiment.

### siRNA transfections

To deplete endogenous p53 for telomere FISH on metaphase spreads experiments, a single siRNA targeting the human TP53 gene was synthesized and purchased from Ambion (Life Technologies). Briefly, 350,000 cells were seeded per well of a 10-cm dish containing growth medium without antibiotics. The following day cells were transfected with Lipofectamine™ RNAiMAX Transfection Reagent (Thermo Fisher). siRNAs and RNAiMAX were diluted in OptiMEM. A working siRNA concentration of 12.5 nM and 6 μl RNAiMAX transfection reagent per 10-cm plate was used. Transfection media was replaced with complete culture media 24 h later. Cells were treated with 20 min DL after 24h and collected after additional 24h. Fot the transfection of OGG1 and Ape1 siRNAs, the same protocol was used with the exception that 30,000 cells were seeded per well of a 35-mm dish and a working siRNA concentration of 5 nM and 3 μl RNAiMAX transfection reagent was used.

### Telomere FISH on metaphase Spreads

Chromosome spreads were prepared by incubating cells with 0.05 μg/ml colcemid for 2 h prior to harvesting with trypsin. Cells were incubated with 75mM KCl for 13 minutes at 37°C and fixed in methanol and glacial acetic acid (3:1). Cells were dropped on washed slides and dried overnight before fixation in 4% formaldehyde. Slides were treated with RNaseA and Pepsin at 37°C, and then dehydrated. FISH was performed as above, and included a CENPB (PNABio) probe in addition to the telomere probe.

### Image Acquisition and Analysis

All IF images were acquired on a Nikon Ti inverted fluorescent microscope equipped with an Orca Fusion cMOS camera, or CoolSNAP HQ2 CCD. Z-stacks were acquired for each image and deconvolved using blind, iterative methods with NIS Elements AR software.

For co-localizations, deconvolved images were converted to Max-IPs and converted to a new document. The object counts feature in NIS AR was used to set a threshold for foci that was kept throughout the experiment. The binary function was used to determine the intersections of 2 or 3 channels in defined regions of interest (ROI) (DAPI stained nuclei). For whole nuclei signal intensity, the automated measurements function was used on ROIs.

### Western Blotting

Cells were collected from plates with trypsin, washed with PBS, and then lysed on ice with RIPA buffer (Santa Cruz) supplemented with PMSF (1nM), 1x Roche Protease and Phosphatase Inhibitors, and Benzonase (Sigma E8263; 1:500) for 15 min and then incubated at 37 °C for 10 min, before spinning down at 15,000 rpm for 10 minutes at 4°C. Protein concentrations were determined with the BCA assay (Pierce) and 20-30 μg of protein was electrophoresed on 4-12% (or 12% for OGG1 blot) Bis-Tris gels (Thermo) before transferring to PDVF or nitrocellulose membranes (GE Healthcare). For PAR 10H western blots, cells were directly lysed right after treatment in 4× LDS sample buffer completed with 20 μM PARPi and 20 μM PARGi. Proteins were denatured for 5 min at 95 °C to inactivate PARP and PARG enzymes. Samples were then gently homogenized using universal nuclease (Pierce/ThermoFisher), resolved by SDS–PAGE electrophoresis and transferred on nitrocellulose. Red Ponceau staining was performed to ensure even transfer of proteins onto membranes, which were then washed in TBS-T and blocked in 5% milk, or 5% BSA for phosphorylated proteins, and blotted with primary and secondary HRP antibodies. Signal was detected by ECL detection and X-ray film.

### Flow Cytometry

To analyze apoptosis, cells were treated as indicated, and allowed to recover for 4 days. Floating cells were collected, and then attached cells were collected with trypsin and combined. After spinning down and washing, cells were incubated with Alexa Flour 488 annexin V and 1 µg/ml propidium iodine in 1x annexin-binding buffer for 15 min in the dark (Thermo). After resuspending in additional binding buffer, cells were analyzed on a CytoFLEX S Flow Cytometer (Beckman) using FL1 and FL3. For cell cycle analysis, 23 h after treatment, cells were pulsed with 20 µM EdU (Thermo), and incubated for an additional hour. Cells were collected with trypsin, washed with 1% BSA in PBS, and then fixed with Click-IT fixative D. After washing with 1% BSA in PBS, the cells were permeabilized with 1x component E for 15 min, before performing Click chemistry with Alexa Flour 488 azide for 30 minutes in the dark. Cells were washed with 1x component E, and then resuspended in 500 µl FxCycle PI/RNase (Thermo) for 15 min before analyzing on Beckman CytoFLEX S Flow Cytometer. Preliminary standard gating for cells versus debris and singlet, and analysis of the results, were conducted with FlowJo™ v10.8 Software (BD Life Sciences).

### Synthesis of OGG1i (TH5487) and OGG1i^NA^

4-(4-Bromo-2-oxo-2,3-dihydro-1*H*-benzo[*d*]imidazol-1-yl)-*N*-(4-iodophenyl)piperidine-1-carboxamide (TH5487 (OGG1i)). This compound was prepared according to literature data^35^ using the following procedures. A mixture of 4-aminopiperidine-1-carboxylic acid *tert*-butyl ester (1.5 g, 7.5 mmol), 1-bromo-3-fluoro-2-nitro-benzene (1.5 g, 6.8 mmol), and diisopropylethylamine (1.7 mL, 10.2 mmol) was stirred in a sealed vial at 120 °C for 16 h. The mixture was concentrated and purified by chromatography on SiO_2_ (50-100% CH_2_Cl_2_ in hexanes) to afford *tert*-butyl 4-((3-bromo-2-nitrophenyl)amino)piperidine-1-carboxylate (2.27 g, 83%) as an orange colored foam: IR (ATR, neat) *v_max_* 3407, 2976, 2932, 2864, 1684, 1598, 1563, 1532, 1497, 1478, 1451, 1424, 1365, 1276, 1239, 1167, 1140, 1094, 1056 cm^-1^; ^1^H NMR (600 MHz, CDCl_3_) δ 7.14 (t, *J* = 8.1 Hz, 1 H), 6.94 (d, *J* = 7.8 Hz, 1 H), 6.76 (d, *J* = 8.4 Hz, 1 H), 5.60 (d, *J* = 6.6 Hz, 1 H), 4.00 (bs, 2 H), 3.54-3.50 (m, 1 H), 2.98 (m, 2 H), 2.00 (d, *J* = 10.8 Hz, 2 H), 1.46 (s, 9 H), 1.44-1.40 (m, 2 H); ^13^C NMR (150 MHz, CDCl_3_) δ 154.7, 142.3, 136.9, 133.0, 121.9, 116.6, 112.8, 80.0, 50.0, 31.8, 28.5; HRMS (ESI^+^) *m/z* for C_16_H_23_O_4_N_3_Br, [M+H]^+^: calcd 400.0867, found 400.0871.

To a stirred solution of *tert*-butyl 4-((3-bromo-2-nitrophenyl)amino)piperidine-1-carboxylate (2.25 g, 5.62 mmol) in acetonitrile (24 mL) and water (2.4 mL) was added NiCl_2_ (0.146 g. 1.12 mmol) at room temperature. The reaction mixture was cooled to 0 °C and NaBH_4_ (0.85 g, 22.5 mmol) was added portionwise over ca. 10 min. The mixture was diluted with CH_2_Cl_2_ (30 mL), decanted, poured into NaHCO_3_ (30 mL) and extracted with CH_2_Cl_2_ (3 × 40 mL). The combined organic layers were dried (Na_2_SO_4_), filtered, and concentrated. The crude residue was purified by chromatography on SiO_2_ (10%-20% EtOAc in hexanes) to afford *tert*-butyl 4-(2-amino-3-bromoanilino)piperidine-1-carboxylate (1.94 g, 93%) as a yellow oily foam: IR (ATR, neat) *v_max_* 3389, 2975, 2932, 2856, 1672, 1585, 1452, 1426, 1392, 1366, 1344, 1311, 1275, 1239, 1218, 1167, 1138, 1069 cm^-1^; ^1^H NMR (400 MHz, CDCl_3_) δ 6.93 (dd, *J* = 1.6, 7.6 Hz, 1 H), 6.63 (t, *J* = 7.8 Hz, 1 H), 6.59 (dd, *J* = 1.6, 8.0 Hz, 1 H), 4.02 (bs, 2 H), 3.79 (bs, 1 H), 3.39-3.32 (m, 2 H), 2.92 (app t, *J* = 11.8 Hz, 2 H), 2.01-1.97 (m, 2 H), 1.46 (s, 9 H), 1.41-1.30 (m, 2 H); ^13^C NMR (100 MHz, CDCl_3_) δ 154.8, 136.6, 133.3, 122.4, 120.5, 112.2, 111.5, 79.6, 50.3, 32.3, 28.5; HRMS (ESI^+^) *m/z* for C_16_H_23_O_4_N_3_Br, [M+H]^+^: calcd 370.1125, found 370.1125.

A mixture of *tert*-butyl 4-(2-amino-3-bromoanilino)piperidine-1-carboxylate (1.94 g, 5.24 mmol), and diisopropylethylamine (2.6 mL, 15.7 mmol) was stirred in CH_2_Cl_2_ (10 mL) at 0 °C. A solution of trichloromethyl carbonochloridate (0.32 mL, 2.6 mmol) in CH_2_Cl_2_ (30 ml) was added dropwise. After 30 min, the mixture was concentrated and purified by chromatography on SiO_2_ (0%-6% MeOH in CH_2_Cl_2_) to give *tert*-butyl 4-(4-bromo-2-oxo-2,3-dihydro-1*H*-benzo[*d*]imidazol-1-yl)piperidine-1-carboxylate (1.95 g, 94%) as a slightly brown foam: IR (ATR, neat) *v_max_* 3158, 2975, 2932, 2864, 1693, 1622, 1593, 1487, 1461, 1424, 1365, 1321, 1287, 1274, 1243, 11655, 1118 cm^-1^; ^1^H NMR (400 MHz, CDCl_3_) δ 10.39 (s, 1 H), 7.15 (dd, *J* = 0.6, 8.0 Hz, 1 H), 7.04 (d, *J* = 8.0 Hz, 1 H), 6.90 (t, *J* = 8.0 Hz, 1 H), 4.50-4.42 (m, 1 H), 4.30 (bs, 1 H), 2.90-2.78 (m, 2 H), 2.32 (d, *J* = 4.4, 8.8 Hz, 1 H), 2.28 (d, *J* = 4.4, 8.8 Hz, 1 H), 1.81 (dd, *J* = 10.4 Hz, 2 H), 1.48 (s, 9 H); ^13^C NMR (100 MHz, CDCl_3_) δ 154.7, 154.5, 129.7, 128.0, 124.0, 122.2, 108.2, 102.8, 80.0, 51.1, 29.2, 28.5; HRMS (ESI^+^) *m/z* for C_16_H_23_O_4_N_3_Br, [M+H]^+^: calcd 396.0917, found 396.0913.

To a stirred solution of *tert*-butyl 4-(4-bromo-2-oxo-2,3-dihydro-1*H*-benzo[*d*]imidazol-1-yl)piperidine-1-carboxylate (1.95 g, 4.92 mmol) in CH_2_Cl_2_ (150 mL) was added TFA (12 mL) at 0 °C. The reaction mixture was warmed to room temperature, stirred for 2 h, concentrated under reduced pressure and dried *in vacuo*. The crude residue was used in the next step without further purification.

A solution of crude residue (0.285 g) from the previous step in CH_2_Cl_2_ (10 mL) was treated with diisopropylethylamine (0.17 mL, 1.0 mmol) and stirred in a sealed tube at room temperature for 5 min. A solution of 4-iodophenylisocyanate (0.14 g, 0.58 mmol) in CH_2_Cl_2_ (5 mL) was added and the mixture was heated at 50 °C for 30 min, cooled to room temperature, and stirred for 2 h. The precipitate was collected by filtration and was washed sequentially with CH_2_Cl_2_, water, MeOH, and CH_2_Cl_2_. The solid was then suspended in boiling MeOH and filtered to give 4-(4-bromo-2-oxo-2,3-dihydro-1*H*-benzo[*d*]imidazol-1-yl)-*N*-(4-iodophenyl)piperidine-1-carboxamide (0.22 g, 80%) as a cream-white solid: IR (ATR, neat) *v_max_* 3261, 2847, 1707, 1638, 1584, 1516, 1485, 1459, 1424, 1372, 1324, 1311, 1300, 1280, 1242, 1157, 1109 cm^-1^; ^1^H NMR (400 MHz, DMSO-d_6_) δ 11.31 (s, 1 H), 8.69 (s, 1 H), 7.57-7.55 (m, 2 H), 7.37-7.35 (m, 2 H), 7.25 (d, *J* = 7.8 Hz, 1 H), 7.16 (d, *J* = 7.8 Hz, 1 H), 6.95 (t, *J* = 8.1 Hz, 1 H), 4.42-4.37 (m, 1 H), 4.28 (d, *J* = 13.2 Hz, 2 H), 2.93 (app t, *J* = 12.0 Hz, 2 H), 2.28 (dd, *J* = 4.2, 12.6 Hz, 1 H), 2.24 (d, *J* =4.2, 12.6 Hz, 1 H), 1.74 (app dd, *J* = 1.8, 10.2 Hz, 2 H); ^13^C NMR (150 MHz, DMSO-d_6_) δ 154.4, 153.4, 140.7, 136.9, 130.3, 127.8, 123.3, 121.9, 121.7, 107.7, 101.1, 84.6, 50.5, 43.4, 28.6.

*N*-(4-chlorophenyl)-4-(2-oxo-2,3-dihydro-1*H*-benzo[*d*]imidazol-1-yl)piperidine-1-carboxamide (OGG1i^NA^). This compound can be obtained analogously according to the experimental protocol provided for OGG1i. The purity of final products was assessed using an Agilent Technologies 1260 Infinity II LC at 220 nm UV absorption (Waters XBridge BEH C_18_ 2.1 × 50 mm, 2.5 µm) or an Agilent Technologies 385-ELSD (Microsolv Cogent 2.0 Bidentate C_18_ 2.1 × 50 mm, 2.2 µm; ELSD conditions: evaporator and nebulizer set at 45 °C; gas flow set at 1.80 standard liter/min). All final assay samples showed a purity >95% by LCMS analysis with UV (220 nm) and ELS detection.

### Statistics and reproducibility

The number of biological and technical replicates are noted in all figure legends and methods. All statistical analysis was perfomed in Graphpad Prism 9. No statistical method was used to predetermine sample size. Investigators were not blinded to allocation during experiments and outcome assessments.

### List of reagents Chemicals

4-(4-Bromo-2-oxo-2,3-dihydro-1*H*-benzo[*d*]imidazol-1-yl)-*N*-(4-iodophenyl)piperidine-1-carboxamide (TH5487 (OGG1i)). See methods.

*N*-(4-chlorophenyl)-4-(2-oxo-2,3-dihydro-1*H*-benzo[*d*]imidazol-1-yl)piperidine-1-carboxamide (OGG1i^NA^). See methods.

Bis[4-(dimethylamino)phenyl](4-(3-carboethoxypropyl)-3,5-diiodo-phenyl)-chloride (MG-2I). prepared as described in ^15, 34^

PDD00017272 – Tocris - Cat#7006; CAS: 1945950-20-8

AZD2281 (Olaparib) - Selleck chemicals – Cat#S1060; CAS: 763113-22-0

### Miscellaneous

Lipofectamine™ RNAiMAX Transfection Reagent - Thermo Fisher – Cat#13778075 ProLong™ Diamond Antifade Mountant - Thermo Fisher – Cat#P36965

### Antibodies

GAPDH (Santa Cruz sc-47724; 1:5000 WB), α-tubulin (Millipore 05-829; 1:5000 WB), OGG1 (Abcam ab124741; 1:500), MUTYH (Abnova H00004595-M01; 1:500 WB); γH2AX (Santa Cruz sc-517348; 1:200 IF), 53BP1 (Novus NB100-304; 1:1000 IF), p53 (Santa Cruz sc-126; 1:200 WB), p21 (Cell Signaling 2947; 1:1000 IF and 1:2000 WB), pCHK2 T68 (Cell Signaling 2197; 1:1000 WB), pCHK1 S345 (Cell Signaling 2348; 1:500 WB), pATM S 1981 (Abcam ab81292; 1:2000 WB), Chk2 (Cell Signaling 3440; 1:1000 WB), CHK1 (Cell Signaling 2360; 1:1000 WB), ATM (Sigma A1106; 1:500 WB); PAR (10H) (Enzo ALX-804-220-R100, 1:500 WB, 1:100 IF), APE1 (Cell Signaling 10519S; 1:1000 WB); Cyclin A (Santa Cruz sc-271682; 1:500 WB).

### Recombinant DNA

pLentiCRISPR v2 gRNA, OGG1 targeting sequence (exon4:GCTACGAGAGTCCTCATATG)

pLentiCRISPR v2 gRNA, MUTYH targeting sequence (exon2:GCATGCTAAGAACAACAGTC)

ON-TARGETplus Non-targeting Pool scramble – Dharmacon - Cat#D-001810-10-05 TP53

siRNA - Ambion (Life Technologies) – Cat#4390824 – sequence (5’ -> 3’) GGUGAACCUUGUACCUAAtt

OGG1 siRNA ON-TARGETplus SMARTPool – Dharmacon - Cat#M-005147-03-0005

APE1 siRNA ON-TARGETplus SMARTPool - Dharmacon – Cat#L-010237-00-0005

## Notes

### Competing Interest Statement

The authors have declared no competing interest.

### Summary of Updates

A different format of the document was uploaded to prevent the bioRXiv stamp (header on top of page) to cover graphs and images in the figure section.

