## Supplementary figures and images for "OGG1 and MUTYH repair activities promote telomeric 8-oxoguanine induced cellular senescence"

### Figure S1

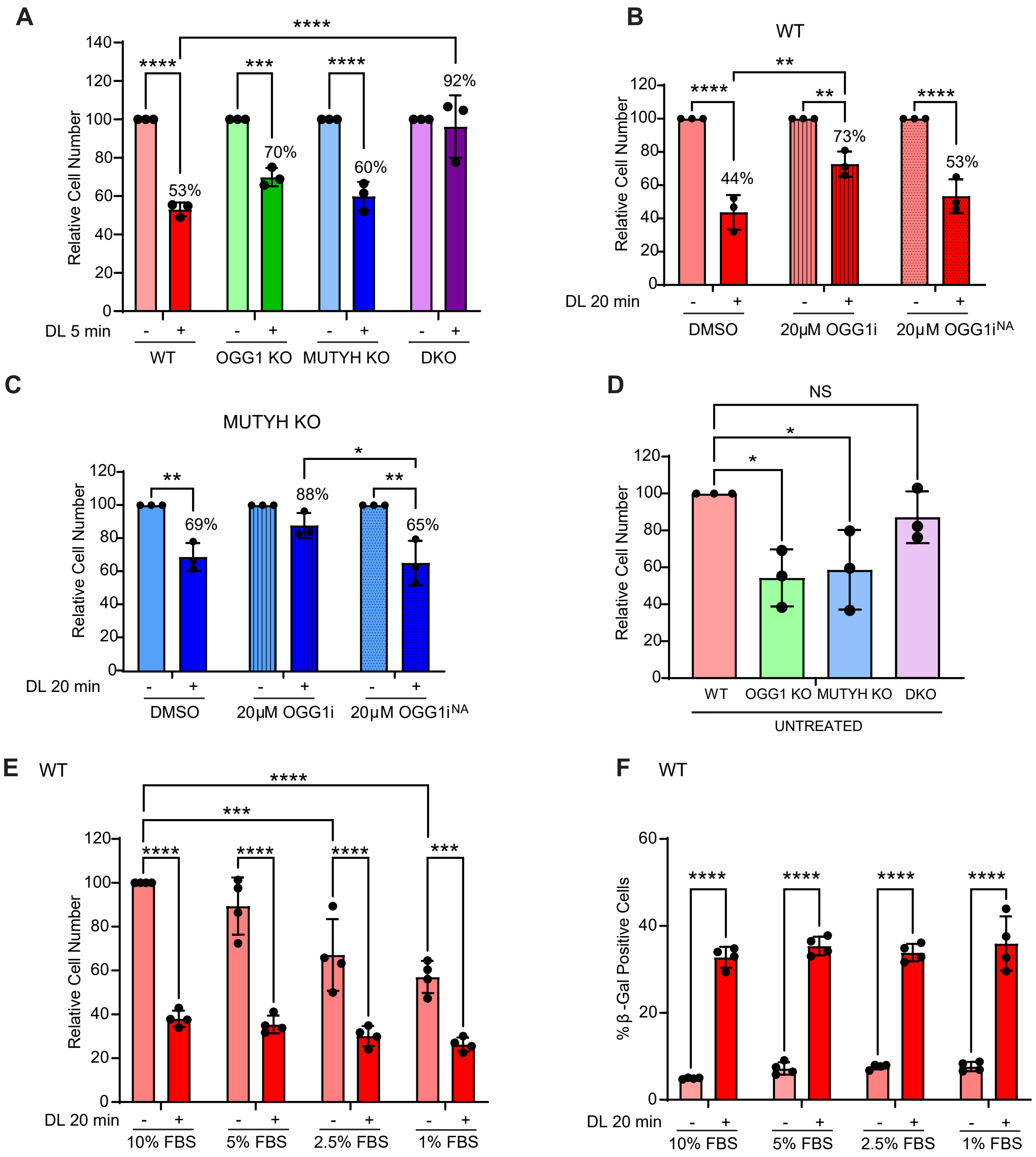

### Figure S2

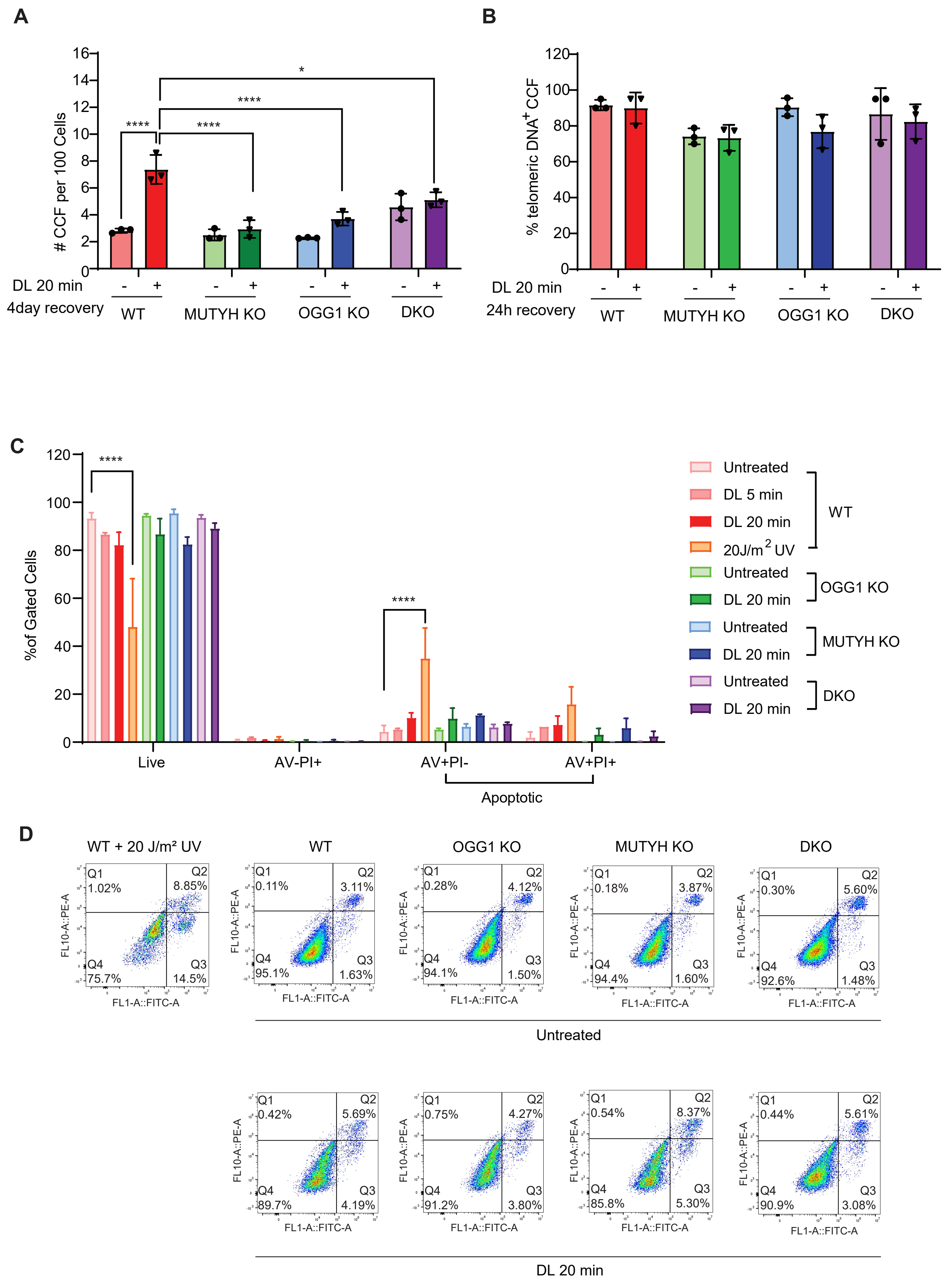

### Figure S3

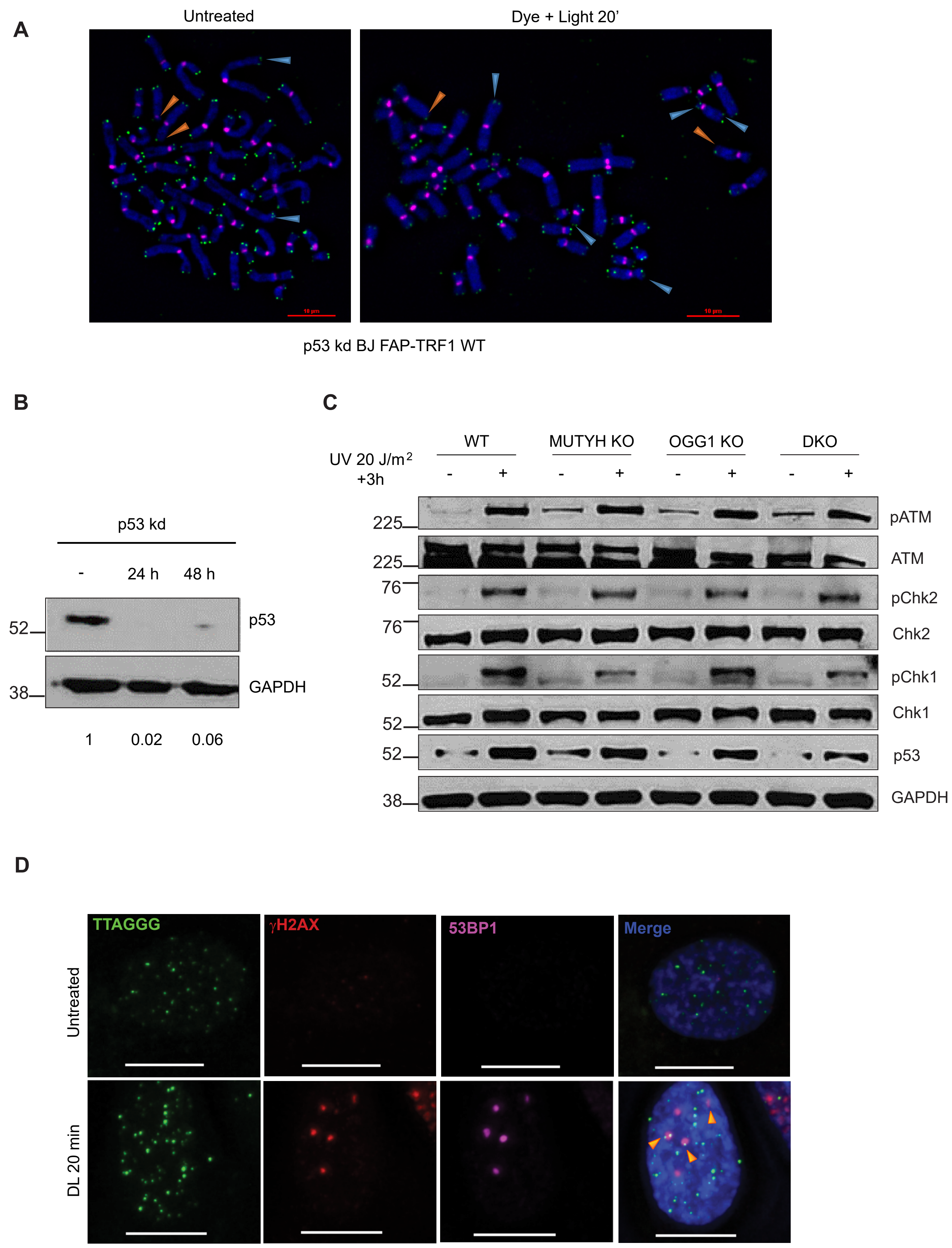

### Figure S4

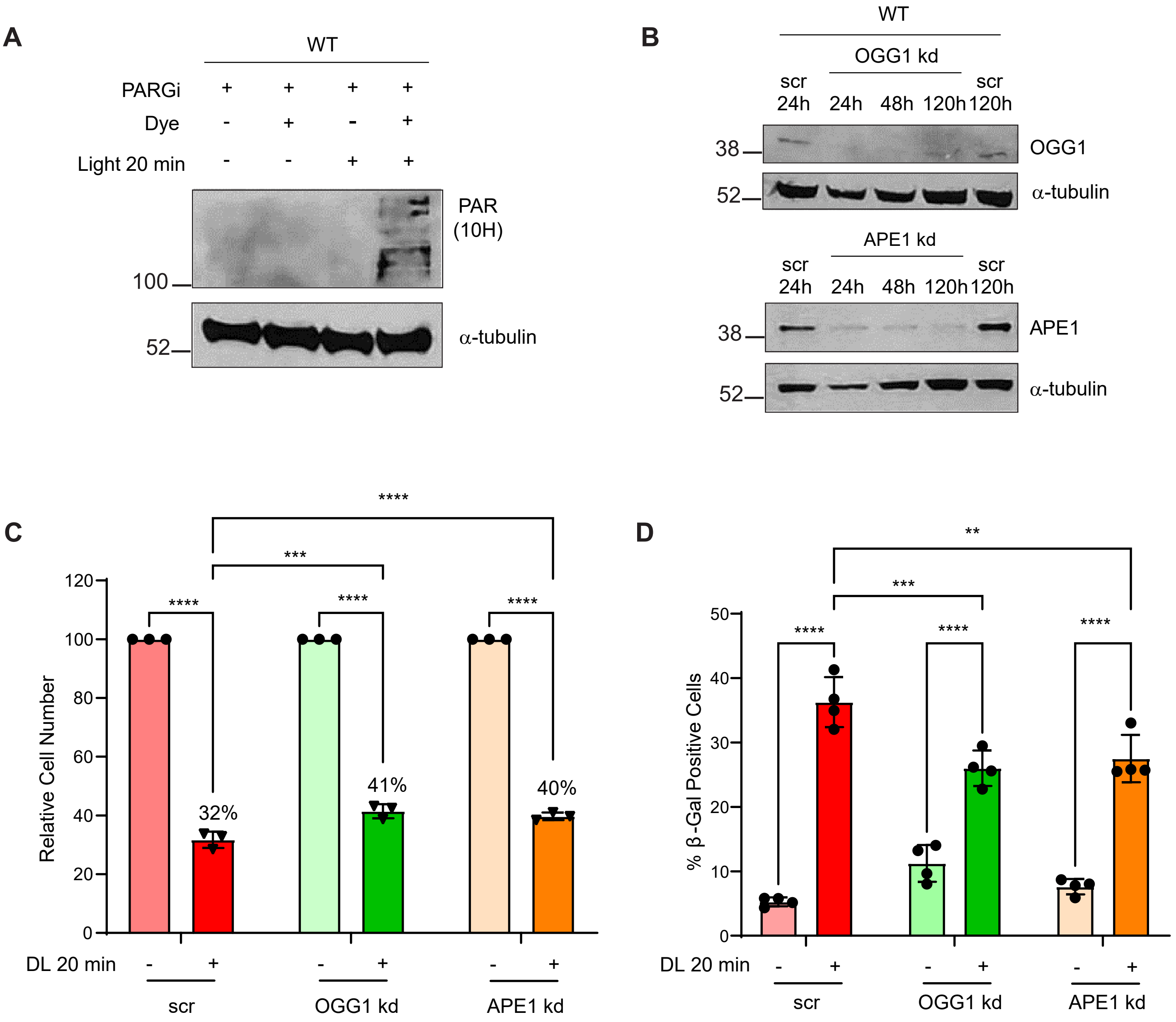

### Figure S5

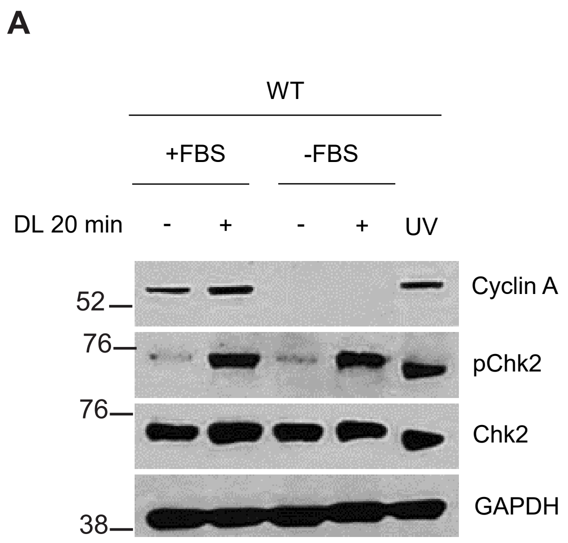

### Figure S6

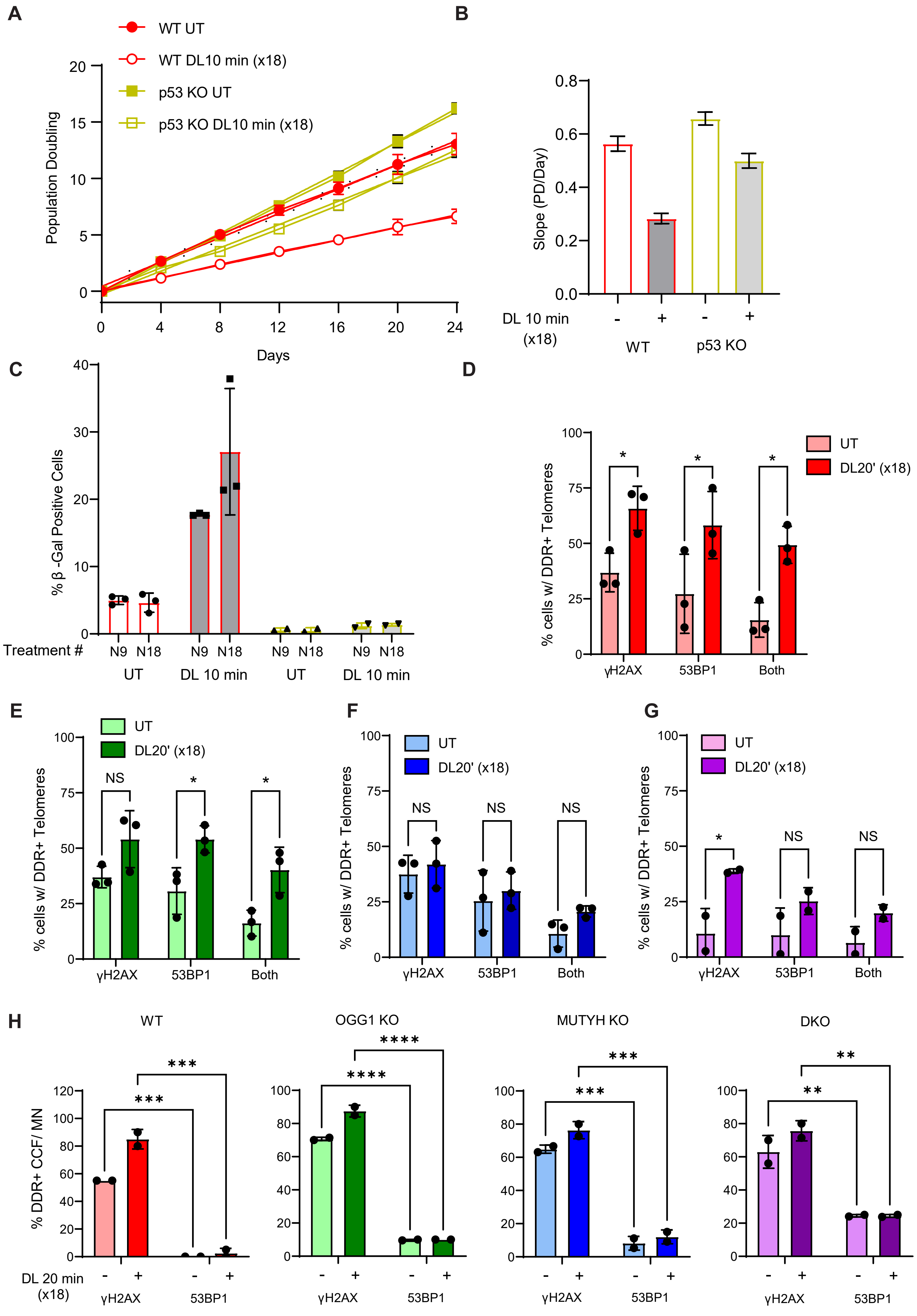
